# Eco-Evolutionary Dynamics of Proliferation Heterogeneity: A Phenotype-Structured Model for Tumor Growth and Treatment Response

**DOI:** 10.64898/2026.03.13.711687

**Authors:** Lara Schmalenstroer, Russell C. Rockne, Farnoush Farahpour

## Abstract

Intra-tumor heterogeneity in proliferation rates fundamentally influences cancer progression and treatment resistance. To investigate how continuous phenotypic variation shapes eco-evolutionary dynamics, we develop a phenotype-structured partial differential equation framework that explicitly models proliferation heterogeneity as a dynamic trait. Our model integrates three key biological principles: (1) phenotypic diffusion capturing heritable variation in proliferation rates, (2) global resource competition enforcing density-dependent growth constraints, and (3) an experimentally grounded life-history trade-off linking elevated proliferation to increased mortality. Using adaptive dynamics, we derive the optimum proliferation rate in a growing tumor, showing that the optimal phenotype dynamically shifts toward slower proliferation as tumors approach carrying capacity under control condition. We perform *in silico* treatment simulations for four different treatment regimes (pan-proliferation, low-, mid-, and high-proliferation targeting) to show how therapeutic selective pressures reshape fitness landscapes. While all treatments slow down tumor growth, they induce divergent evolutionary trajectories. We connect these dynamics with changes in mean proliferation rates during and after treatment. Our work establishes a predictive, evolutionarily grounded framework for understanding how therapy reshapes tumor proliferation landscapes, offering a mechanistic basis for designing strategies that anticipate and counteract adaptive resistance.

## 1 Introduction

Biological populations are inherently heterogeneous. Even genetically related individuals can differ substantially in phenotypic traits such as reproduction rate (Travers et al. 2015), metabolic activity (Zhang et al. 2025), and stress tolerance (King et al. 2004). This diversity arises through genetic recombination, epigenetic plasticity (Richards 2006), and stochastic gene expression (Raj and van Oudenaarden 2008), thereby creating complex evolutionary landscapes where selection, mutation, and environmental pressures drive dynamic adaptation (de Visser and Krug 2014). Crucially, phenotypic heterogeneity and variation operate within fundamental biophysical constraints: organisms cannot optimize all phenotypic traits simultaneously due to finite energetic resources (Shoval et al. 2012). These universal fitness trade-offs (Stearns 1989) manifest across biological scales. For example, in animals, rapid reproduction often comes at the cost of reduced longevity (Ghalambor and Martin 2001; Partridge et al. 2005; Travers et al. 2015; Lemaître et al. 2015), while in microbes, slow growth frequently enhances stress resistance (King et al. 2004; Lu et al. 2009; Zakrzewska et al. 2011; Phan and Ferenci 2013). More recently, human cell studies have shown that genomic variants favoring proliferation can decrease survival under anticancer therapies (Kim et al. 2024), demonstrating how these principles extend to pathological contexts. Analogous compromises govern immune function (McDade et al. 2016; Schwenke et al. 2016), metabolic specialization (Ganeshan et al. 2019; Ekkers et al. 2022), and drug tolerance (Wood and Cluzel 2012; Schlatter and Kinkel 2015; Herren and Baym 2022).

In cancer, these universal biological principles manifest as intra-tumor heterogeneity (ITH) (Heppner 1984), which represents a fundamental driver of disease progression (Caswell and Swanton 2017) and therapeutic resistance (Maley et al. 2017; Marusyk et al. 2020; Vitale et al. 2021), and can also influence metastasis (Sobral et al. 2022; Lucas et al. 2025). Within a single tumor mass, cancer cells exhibit marked variability in their traits, including proliferative (Lucas et al. 2025) and metabolic capacity (Meng et al. 2024; Kondo et al. 2021), and treatment sensitivity (Marusyk et al. 2020). This diversity creates complex evolutionary landscapes where subpopulations compete and adapt under selective pressures. Central to this variability is the dynamic heterogeneity in cellular proliferative capacity (Chu et al. 2004; Oren et al. 2021), a key determinant of tumor growth kinetics and fitness, that facilitates the exploration of alternative adaptive strategies by tumor cells under distinct conditions, such as untreated expansion or therapeutic stress, each of which may selectively favor specific proliferative ranges.

Various mathematical approaches have been used to model proliferation heterogeneity in cancer. Compartmental ordinary differential equations (ODE) models discretize populations into subpopulations with distinct proliferative capacities, allowing for mechanistic investigation of selection, treatment response, and clonal competition under therapy (Raatz et al. 2021). Agent-based models (ABMs) (Macal and North 2005) excel at simulating emergent spatial and phenotypic heterogeneity through stochastic cell-level behaviors (Farahpour et al. 2018) and are widely used in computational oncology (West et al. 2023). In particular, ABMs have been employed to investigate evolutionary trade-offs between proliferation and migration rates (Gallaher et al. 2019) as well as proliferation rate and treatment resistance (Gatenby et al. 2020) and their role in shaping phenotypic evolution (Farahpour et al. 2018). Stochastic formulations provide complementary approaches for representing proliferation rate heterogeneity through probabilistic dynamics (Raatz et al. 2021; Durrett et al. 2011; Wang et al. 2023). Adaptive dynamics provides a theoretical framework for modeling long-term evolution under frequency-dependent selection. Foundational models in the field have investigated the evolutionary trajectories of systems characterized by continuous adaptive traits, such as proliferation, migration or therapy resistance, explicitly coupling ecological interactions with evolutionary change, emphasizing the critical role of life-history trade-offs (Brännström et al. 2013; Dieckmann and Law 1996). Finally, phenotype-structured PDE frameworks enable the continuous tracking of evolving trait distributions at the population level. Such trait-structured models have been employed to investigate phenotypic characteristics, including cell motility (Fiandaca et al. 2022) and drug resistance (Lorenzi et al. 2016; Cho and Levy 2018). However, to the best of our knowledge, the proliferation rate has not yet been formulated as an explicit evolving phenotypic trait within this framework.

To investigate the direct impact of evolving proliferative capacity, we formulate a proliferation-structured PDE framework that integrates intra-tumor heterogeneity with a classical tumor growth model while enabling evolutionary analysis. By treating proliferation as a continuously evolving phenotypic trait rather than a fixed parameter, our model provides a mechanistic description of how heterogeneity in growth contributes to macroscopic tumor dynamics. This formulation allows us to relax the conventional assumption of a fixed growth rate, whether at the level of the entire population or of individual subpopulations, and to assess its consequences within a structure that remains directly comparable to fundamental population growth models. Our model integrates three components critical for tumor growth modeling: (1) Phenotypic diffusion captures stochastic heritable variation driving ITH, a fundamental feature of cancer progression and therapeutic resistance; (2) Global resource competition models systemic nutrient limitations observed in solid tumors, where total tumor burden constrains growth uniformly across all phenotypes; and (3) Explicit life-history trade-offs quantify the fitness cost of proliferation strategies observed in oncology. Within our PDE framework, validation against longitudinal tumor volume data demonstrates that the life-history trade-off is essential for accurate growth heterogeneity modeling, preventing unrealistic inflation of mean proliferation rates. Using principles from adaptive dynamics (Brännström et al. 2013), we derive the maximum fitness strategy for proliferation rates under resource competition and show how this maximum dynamically shifts during tumor progression, revealing how tumors restructure phenotypic landscapes during growth and treatment response.

The model also provides a simple framework to investigate how selective pressures acting on specific ranges of proliferation rates shape the eco-evolutionary dynamics of the population. Different anticancer therapies impose different selective pressures across the proliferation spectrum. Conventional cytotoxic chemotherapy is classically most effective against rapidly dividing cells because many agents disrupt DNA synthesis or mitosis, which makes rapidly proliferating tumor subpopulations particularly vulnerable (Raatz et al. 2021). Radiotherapy also tends to preferentially damage actively cycling tumor cells, although its efficacy is additionally shaped by cell-cycle phase, DNA-repair capacity, and microenvironmental factors such as hypoxia (Baskar et al. 2014). By contrast, immunotherapy is not primarily coupled to a tumor cell’s proliferation rate, instead, it works by activating or releasing immune effector mechanisms against antigen-bearing cancer cells, so its activity is generally less selective for rapidly proliferating phenotypes than chemo- or radiotherapy. Through *in silico* treatment experiments, the model demonstrates how strategies that target cells with different proliferative capacities drive distinct evolutionary trajectories of the population. By integrating concepts from ecology, evolution and cancer biology, this framework provides a predictive platform for probing treatment-induced selection and resistance evolution.

## 2 Mathematical Model

We introduce a phenomenological phenotype-structured PDE model to describe heterogeneous tumor growth with continuous variability in cellular proliferative capacity. The model resolves tumor cell populations in time *t* and in a continuous phenotypic trait *ρ*, which represents the intrinsic proliferation rate of individual cells. Two coupled population densities are considered: viable/proliferative cancer cells (*C*_*v*_(*t, ρ*)) and doomed cells (*C*_*d*_(*t, ρ*)). For each time point *t*, the functions *C*_*v*_ and *C*_*d*_ quantify how cells are distributed across the phenotypic axis *ρ*, thereby capturing intra-tumoral heterogeneity in proliferation rates. Proliferation occurs exclusively in viable cells and follows logistic growth kinetics to capture density-dependent resource limitations, while doomed cells undergo no division and will eventually be cleared from the system. We impose a life-history trade-off where elevated proliferation rates correlate with reduced cellular longevity due to resource allocation conflicts. Stochastic variation in the heritable phenotypic trait is modeled as diffusion across the proliferation rate spectrum.

The phenotypic dynamics governing ITH in proliferation rates are quantified by a coupled PDE system (Eq. 1a-g):

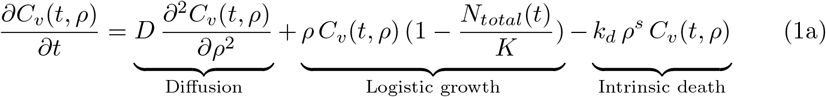

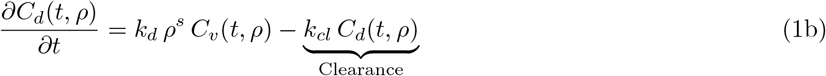

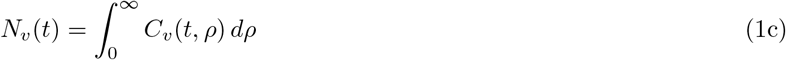

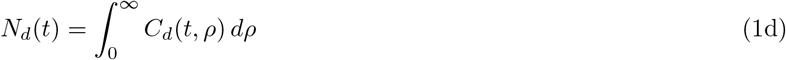

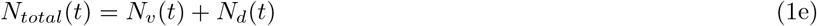

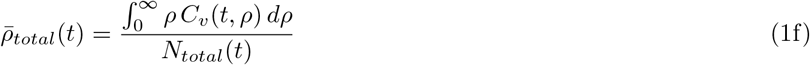

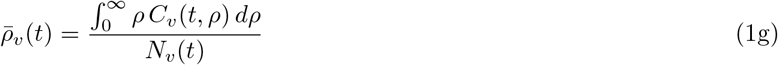

Equation (1a) describes the dynamics of viable cancer cells *C*_*v*_(*t, ρ*), incorporating three fundamental processes: (1) The diffusion term formalizes stochastic heritable variation in proliferation phenotypes, capturing genetic and epigenetic instability, driven, for example, by replication errors, epigenetic fluctuations, or unbiased mutations in regulatory genes. This density-independent formulation models unbiased phenotypic exploration without implying spatial movement. By construction, this diffusion term describes incremental, continuous changes in proliferation rates, and thus cannot represent rare but large or discontinuous phenotypic shifts. (2) The logistic growth term models resource-limited proliferation, where *K* denotes the environment’s carrying capacity and *N*_*total*_(*t*) is the total tumor volume. The density-dependent regulation is assumed to depend on the total tumor burden, leading to a nonlocal uniform coupling across phenotype space. This formulation enforces global competition for shared resources while preserving phenotype-specific intrinsic fitness differences. Consequently, density dependence controls ecological saturation through global regulation rather than phenotype-specific competitive coefficients. This is biologically plausible in the absence of mechanisms linking proliferation rate to resource specialization or spatial niche segregation. (3) The death term *k*_*d*_*ρ*^*s*^*C*_*v*_(*t, ρ*) models irreversible transitions from viable to doomed state. We implement a life-history trade-off, with *k*_*d*_ scaling mortality intensity and *s >* 0 determining the shape of the proliferation-dependent penalty. This trade-off reflects increased mortality risks for rapidly proliferating phenotypes due to mechanisms like metabolic exhaustion and DNA damage accumulation (Aktipis et al. 2013; Boddy et al. 2018). An alternative form of trade-off is discussed in the supplementary section S2.5. Without a life-history trade-off, the combination of phenotypic diffusion and proliferation-driven growth would lead to a systematic drift toward higher population-wide proliferation rates and unrealistic tumor growth. The trade-off term serves as a stabilizing mechanism, ensuring biologically plausible evolutionary dynamics by counterbalancing the fitness advantage of rapid proliferation (see Section S2.1, Figure S2).

Equation (1b) describes doomed cell dynamics *C*_*d*_(*t, ρ*), where the source term *k*_*d*_*ρ*^*s*^*C*_*v*_(*t, ρ*) couples *C*_*d*_ to the population of viable cells by quantifying transitions from viable to doomed phenotypic states. Accumulation occurs proportionally to viable cell death rates, with clearance governed by the clearance rate *k*_*cl*_. Doomed cells retain the proliferation label inherited at transition but do not proliferate or undergo further phenotypic variation.

Integrating the compartments of viable (*C*_*v*_) and doomed (*C*_*d*_) cells across the proliferation rate spectrum yields the volume of the viable, *N*_*v*_(*t*), and doomed, *N*_*d*_(*t*), compartments, respectively (Eq. (1c), (1d)). The total tumor volume, *N*_*total*_(*t*), is calculated as the sum of the viable and doomed volumes (Eq. (1e)). This global metric quantifies tumor progression by linking microscopic phenotypic heterogeneity to macroscopic tumor size.

The population-wide mean proliferation rate 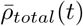 (Eq. (1f)) is defined as the first moment of the viable cell proliferation rate distribution *ρ C*_*v*_(*t, ρ*) *dρ*, normalized by the total tumor volume *N*_*total*_(*t*), quantifying the tumor’s effective proliferative capacity per total tumor burden. The mean proliferation rate of viable cells, 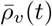, is defined analogously by considering only the volume of the viable cell subpopulation (Eq. (1g)). Details regarding boundary conditions, numerical implementation, parameter calibration, and used data are provided in the supplementary material (see Section S1).

## 3 Results

### 3.1 Model outcome: from macroscopic growth dynamics to microscopic phenotypic restructuring

To characterize the model’s long-term behavior, we simulated the model for 2000 days for a representative tumor with initial and boundary conditions described in Section S1.1. The total tumor volume *N*_*total*_(*t*) initially grows toward carrying capacity but then stabilizes slightly below *K* (Figure 1a). During tumor growth, the viable subpopulation *N*_*v*_ (orange curve in Figure 1b) initially increases, followed by a decline once the tumor approaches carrying capacity. In contrast, the doomed subpopulation *N*_*v*_ (green curve in Figure 1b) increases logistically over time. Both populations reach a steady state in the long run. The persistent volume of non-proliferative cells can represent cell states like necrosis or senescence.

**Fig. 1.**
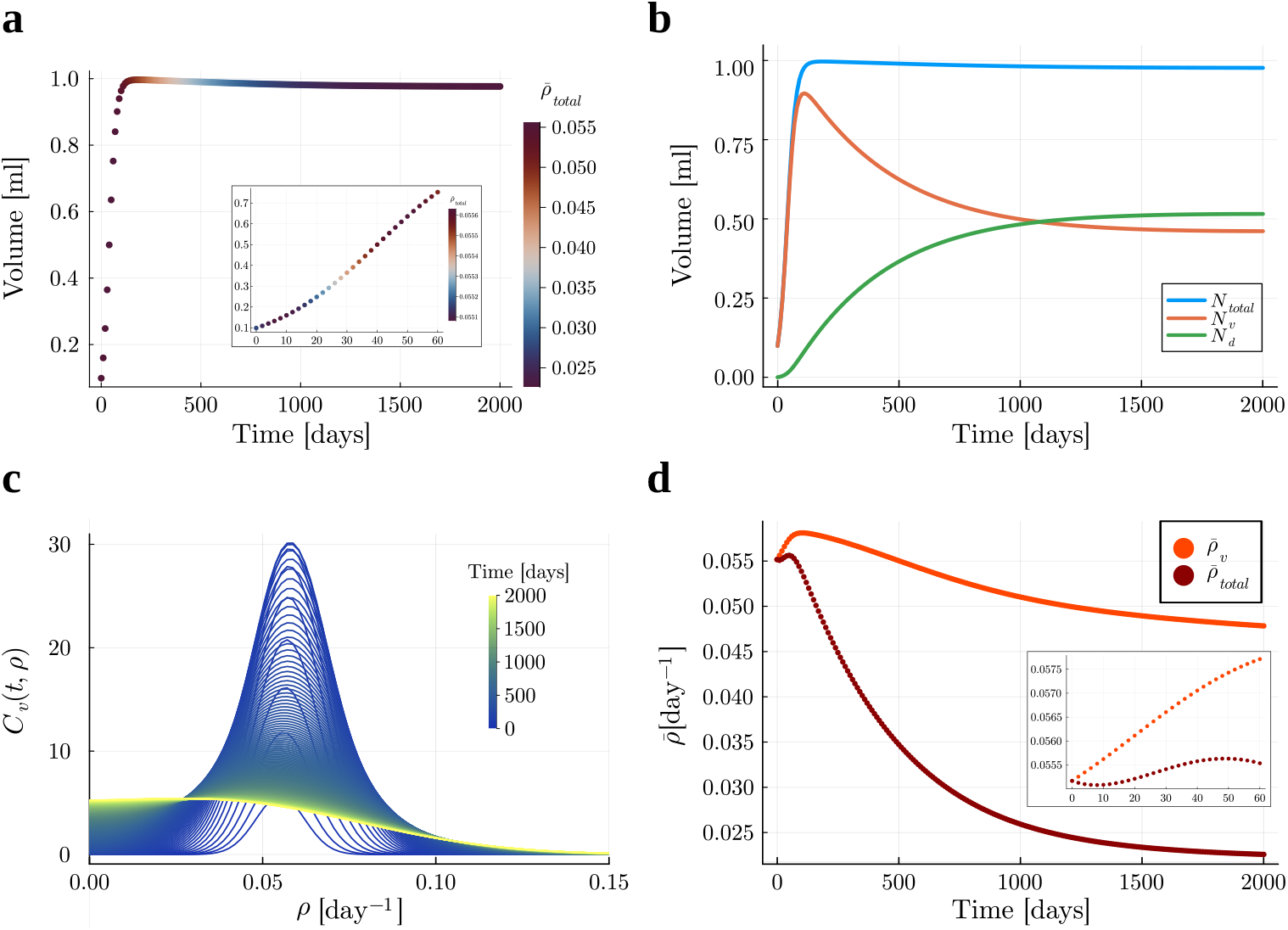
Long-term model behavior for a tumor with the parameters *N*_*total*,0_ = 0.01 [ml], *K* = 1 [ml] *s* = 1.4, *k*_*d*_ = 0.07 [day^(1*−*s)^], *k*_*cl*_ = 0.001 [day^*−*1^], *D* = 5 × 10^*−*7^ [day^*−*3^], *σ*_0_ = 0.007 [day^*−*1^] and 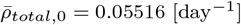 for a time period of 2000 days. **(a)** Tumor volume trajectory: PDE solution for the total tumor volume *N*_*total*_(*t*) (colored dots, hue scaled by population-wide mean proliferation rate 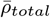). The inset highlights the early-time dynamics **(b)** Trajectories of the total tumor volume *N*_*total*_ (blue curve), the volume of the viable subpopulation *N*_*v*_ (orange curve), and the volume of the doomed subpopulation *N*_*d*_ (green curve). **(c)** Phenotypic restructuring: Viable cell distributions *C*_*v*_(*t, ρ*) from *t*_0_ = 0 days (dark blue) to *t*_*end*_ = 2000 days (yellow). **(d)** Proliferation rate evolution: Temporal dynamics of 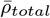 (entire population, red) and 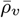 (viable subpopulation, orange). The inset highlights the early-time dynamics.

This phenotypic restructuring is evident in the evolving viable cell distribution *C*_*v*_(*t, ρ*) (Figure 1c): the initial normal distribution (dark blue) centered around the initial mean proliferation rate grows and broadens during early growth, then flattens and skews toward lower proliferation rates as the tumor stabilizes (yellow). This shift arises from two key processes: diffusion spreads the distribution and intrinsic mortality selectively removes rapidly proliferating cells. The resulting decline in population-wide mean proliferation rate 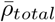 (red, Figure 1d) mirrors the phenotypic shift, initially rising during growth, then falling to a steady state. The viable subpopulation’s mean proliferation rate 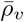 (orange) follows a similar trajectory of initial increase followed by a monotonic reduction, confirming that phenotypic selection occurs within the viable compartment. The final proliferative composition of the tumor reflects the coexistence of non-proliferative cells (e.g., necrotic, senescent, or quiescent states) alongside proliferative cells distributed across a broad spectrum of proliferation rates due to a selection-mutation balance.

### 3.2 Life-history trade-off as evolutionary constraint

To characterize the long-term evolutionary behavior of the proliferation-structured system, we employ the adaptive dynamics formalism to analyze the state-dependent selection gradient and identify evolutionarily singular strategies. In particular, we examine how the proliferation–mortality trade-off term −*k*_*d*_ *ρ*^*s*^ *C*_*v*_ constrains and shapes the selection of optimal proliferative phenotypes. For the proposed trade-off function, any *s >* 0 yields a monotonically increasing mortality rate with respect to *ρ*. When *s >* 1, the trade-off term is convex (Figure 2a, dark blue curve): mortality increases more than proportionally at high proliferation rates. This generates a steep negative selection gradient at large *ρ* and thereby imposes strong evolutionary constraints on further increases in proliferation. For *s* = 1, mortality increases proportionally with proliferation (Figure 2a, mid blue curve), implying a constant selection gradient. For 0 *< s <* 1 (Figure 2a, light blue curve), mortality increases sublinearly with the proliferation rate, thereby reducing the incremental penalty for more rapid proliferation. *s* = 0 represents a simple logistic model with a constant intrinsic death rate, without trade-off.

**Fig. 2.**
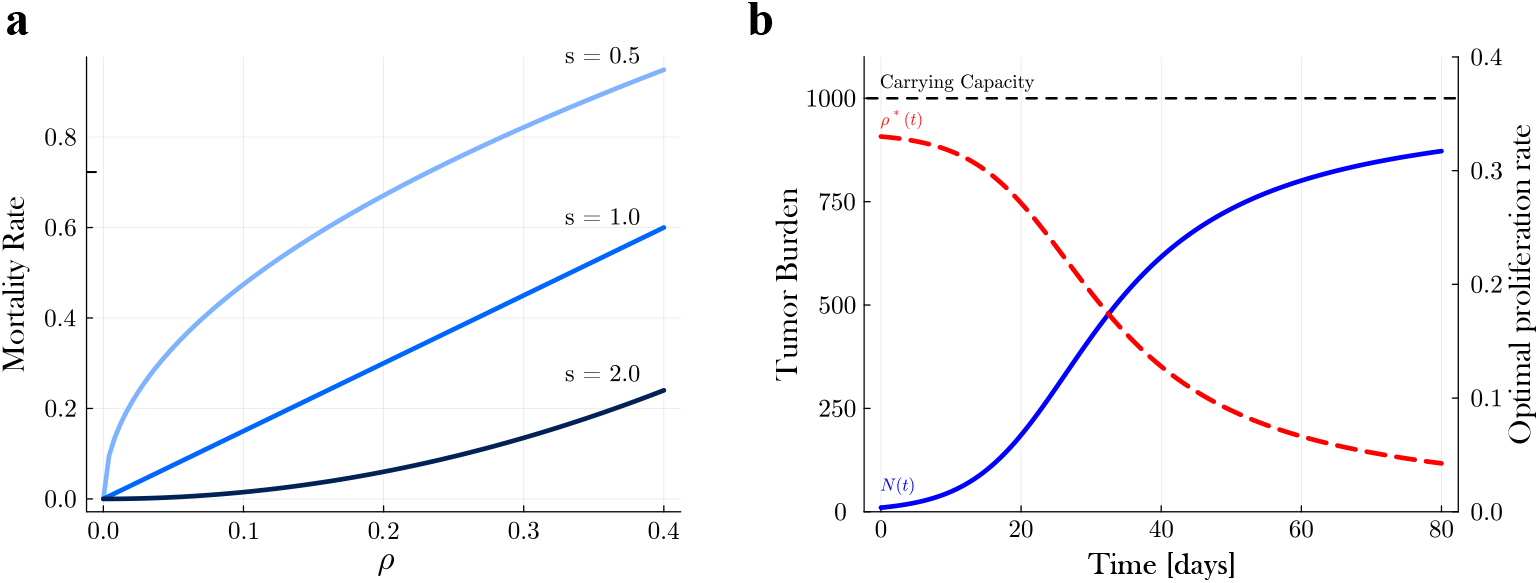
Evolutionarily analysis of tumor growth dynamics based on trade-off shape**(a)** Mortality rate *k*_*d*_*ρ*^*s*^ as a function of proliferation rate *ρ* for trade-off steepness parameters *s* = 0.5 (light blue curve), *s* = 1 (mid blue curve), and *s* = 2 (dark blue curve) and for *k*_*d*_ = 1.5. **(b)** Tumor burden for an exemplary tumor with the parameters *N*_0_ = 10 mm^3^, *K* = 1 mm^3^, *k*_*d*_ = 1.5 days, and *s* = 2.0. Tumor burden *N* (*t*) (blue curve) increases logistically towards carrying capacity *K* (black dashed line). The maximum proliferation rate *ρ*^*^ (red curve) dynamcially decreases as tumor volumes approaches carrying capacity.

This trade-off, counteracting the fitness gain by direct proliferative capacity, reshapes phenotypic fitness, as represented in the net growth rate *g*(*ρ*) which defines a phenotype-dependent fitness landscape governing evolutionary dynamics:

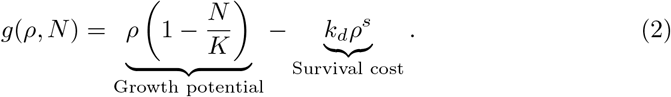

Depending on the value of *s*, this function produces distinct fitness landscapes. An interior fitness maximum exists only for *s >* 1, for which the net growth rate is strictly concave in *ρ*. This concavity condition (∂^2^*g/*∂*ρ*^2^ *<* 0) arises because *∂*^2^*g/∂ρ*^2^ = −*k*_*d*_*s*(*s* − 1)*ρ*^*s−*2^ is strictly negative when *s >* 1. When *s* ≤ 1, no interior maximum fitness exists. At *s* = 1, the second derivative vanishes, while for *s <* 1 it becomes positive, yielding a minimum, forcing the fitness optimum to the boundaries.

To derive the dynamic optimal evolutionary strategy, *ρ*^*^, we set the first derivative of *g*(*ρ, N*) equal to 0, that gives:

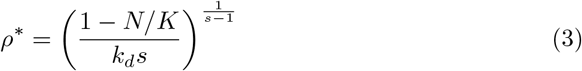

The net growth rate *g*(*ρ*) and its maximizing trait, *ρ*^*^, are dynamically coupled to tumor volume *N* (*t*). This is an example of eco-evolutionary dynamics, in which the environment is shaped by the strategies of all individuals, which in turn changes the optimal evolutionary strategy dynamically. For *s >* 1, as *N* (*t*) (Figure 2b, blue curve) approaches the carrying capacity *K*, the growth potential term 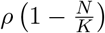 is progres-sively suppressed, leading to a decrease of *ρ*^*^ (red dashed curve) with 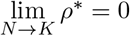. In this ecologically stable state, selection favors vanishing proliferation due to complete resource saturation. In this limit, our PDE with the diffusion term, keeps the system in a selection-mutation balance, where the population maintains a low but non-zero proliferation rate. For *s <* 1, by contrast, 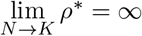, implying that, under any biologically constrained proliferation interval, the evolutionary attractor is driven toward the upper boundary of the admissible trait space. Such behavior corresponds to run-away selection for maximal proliferation and is therefore not biologically plausible. Consequently, we restrict our analysis to trade-off exponents *s* ≥ 1. The value of *s* in this regime strongly influences the proliferation profile and, consequently, tumor volume dynamics. Higher values of *s* impose a greater penalty on rapidly proliferating cells, leading to a reduction in both the overall growth rate and the eventual saturation level of the tumor. From an evolutionary perspective, this suggests that cancer cells may benefit from adapting internal regulatory mechanisms that effectively adjust trade-off shape, thereby optimizing proliferation at the population level.

Further details on the dynamic reshaping of the fitness landscape, including the evolution of the net growth rate *g*(*ρ, N*) are provided in the supplementary material (Section S2.4, Figure S5).

### 3.3 Replicate-specific parameterization of the heterogeneous growth model

The PDE model (Eq. 1a-g) was fitted to longitudinal tumor volume measurements from 11 tumor replicates (see Sections S1.3 and S1.4).

The model solutions are visualized for a representative replicate (Figure 3). The predicted growth trajectory shows close agreement with the experimental measurements (Figure 3a). The model accurately captured the logistic growth pattern, while simultaneously resolving the underlying phenotypic evolution (Figure 3b-c). Fits for the remaining replicates are provided in the supplementary material (see Section S2.3). Temporal patterns in 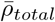 (color-coded in Figure 3a) were observed across the experimental period, with detailed dynamics illustrated in Figure 3b. The mean proliferation rate of the entire population 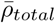 exhibited an early decline, a peak around day 45, and a decrease toward the end of the observation period (red dots). In contrast, the proliferation rate of the viable subpopulation 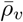 steadily increased over time (orange dots) in this time window.

**Fig. 3.**
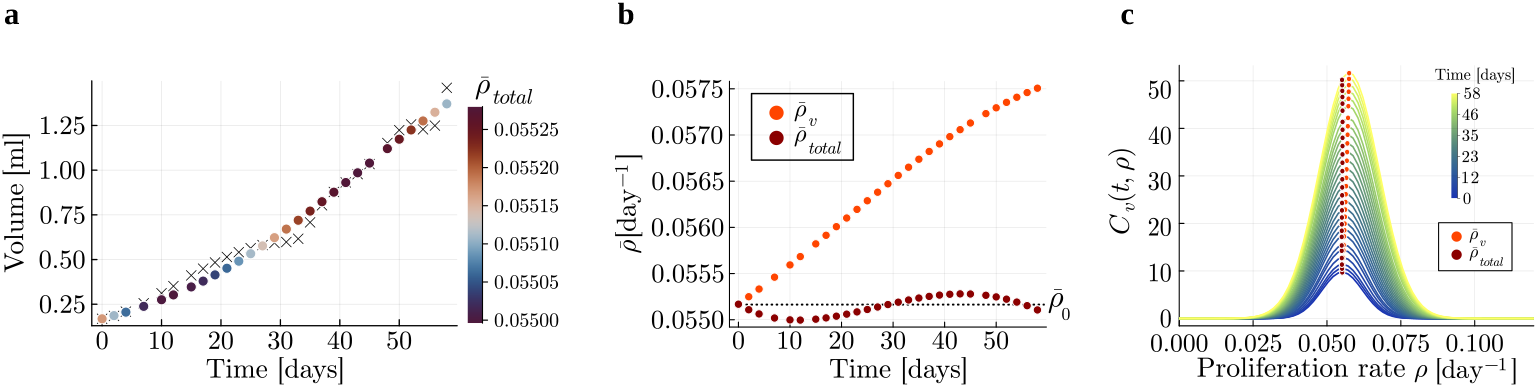
Model validation and phenotypic dynamics for tumor replicate 174 **(a)** Tumor volume trajectory: PDE solutions (colored dots, hue scaled by population-wide mean proliferation rate 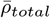) versus experimental measurements (black crosses). **(b)** Proliferation rate evolution: Temporal dynamics of 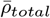 (entire population, red) and 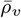 (viable subpopulation, orange). Black dotted line indicates ini-tial 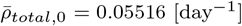. **(c)** Phenotypic restructuring: Viable cell distributions *C*_*v*_(*t, ρ*) from *t*_0_ = 0 days (dark blue) to *t*_*end*_ = 58 days (yellow). Red and orange dots mark 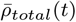 and 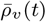 at sampled time points. Parameters *D* = 4.87 × 10^*−*7^ [day^*−*3^], *σ*_0_ = 0.0065 [day^*−*1^], *s* = 3.96, *k*_*d*_ = 97.76 [day^(1*−*s)^], and *k*_*cl*_ = 0.001 [day^*−*1^] were fitted; carrying capacity *K* = 1.97 [ml] was replicate-specific.

A continuous redistribution of the population composition in phenotype occurred throughout tumor progression, as visualized in Figure 3c. The initial Gaussian profile (dark blue) increased in amplitude, while progressively broadening as a consequence of diffusion-driven phenotypic variability, together reflecting tumor growth. Crucially, the mean proliferation rate of the tumor 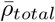 (red dots) systematically resided left-ward of the viable distribution’s mean 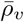 (orange dots) throughout tumor progression. The dynamics in 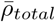 and 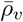 arise from multiple processes: Phenotypic diffusion continuously introduces both slow- and rapid-proliferating cells, while more rapidly proliferating phenotypes intrinsically dominate population growth, increasing both 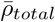 and 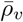. Conversely, the trade-off mortality selectively eliminates rapidly proliferating cells, introducing doomed cancer cells without proliferative capacity into the system, reducing both 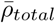 and 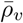. The clearance rate *k*_*cl*_ further modulates this effect by determining the retention time of doomed cells. These mechanisms explain why the most abundant phenotype (distribution peaks) does not align with population-averaged proliferation rates 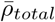 but instead tends to align more closely with the viable population’s mean proliferation rate 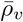. However, due to the asymmetry of the distribution, the peak does not exactly coincide with the viable population’s mean proliferation rate as the shape of the distribution can shift slightly depending on the proliferation-mortality trade-off.

As the tumor approaches carrying capacity, a microenvironment-driven adaptive shift towards slower proliferation occurs, as predicted by the maximum fitness strategy (see Section 3.2). This shift leads to a decrease in 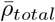 towards the end of the experimental observation window.

The fitted parameters for all replicates can be found in Table S1 in Section S2.3. Figures S3 and S4 show the fitted tumor volumes and the densities over time, respectively, for all fitted replicates. Section S2.5 shows the results of calibration for a model with an alternative trade-off term.

By incorporating heterogeneous proliferation rates into the growth model, our model predicts tumor growth with a dynamically changing tumor composition. This approach allows modeling the evolution of tumor growth and the emergence of sub-populations under different selective pressures (see Section 3.4).

### 3.4 Treatment-driven selective pressures reshape tumor phenotypic landscapes

With the experimental validation of our PDE framework’s tumor growth simulations, we now investigate how cancer therapies reshape tumor dynamics through selective pressures.

To model therapeutic interventions, we extend the system by adding a treatment term −*f* (*ρ*) · *C*_*v*_(*t, ρ*) to the original equations, where *f* (*ρ*) represents the proliferation-dependent killing rate. The extended model becomes:

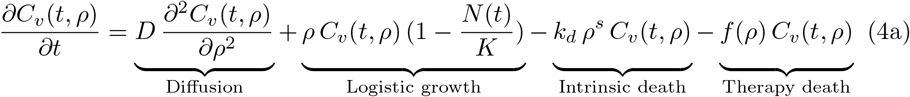

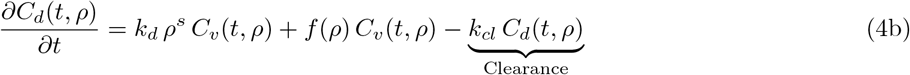

Through *in silico* experiments, conducted without additional parameter fitting, we simulate four mechanistically distinct regimens applied as 10-day continuous treatments initiated at day 5. By comparing responses across targeting strategies, we identify how treatment-driven selection modulates the fitness trade-offs inherent in heterogeneous tumors. The areas under the curve of each treatment profile are normalized to enable comparative analysis of targeting strategy rather than dose intensity.

#### Pan-proliferation targeting

Cells are affected independently of their proliferation rate *ρ*, implementing uniform suppression across the phenotypic spectrum. The constant treatment effect follows *f*_*uniform*_(*ρ*) = *k*, where *k* is a constant value applied identically to all proliferation rates (blue horizontal line in Figure 4a).

**Fig. 4.**
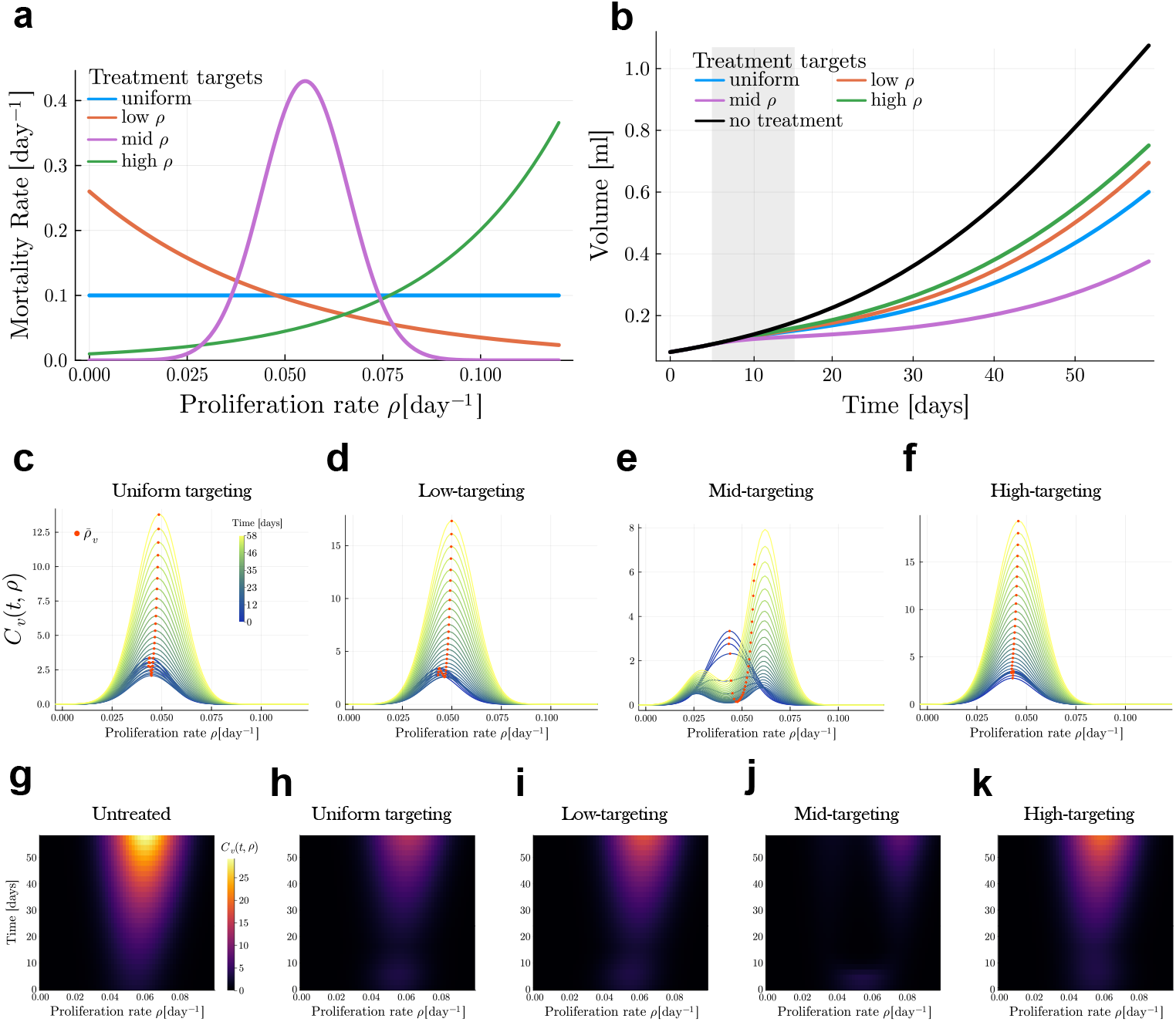
Treatment-induced restructuring of phenotypic landscapes. **(a)** Therapeutic targeting profiles: Suppression functions for uniform (blue), slow-proliferation-targeted (orange), mid-proliferation-targeted (purple), and high-proliferation-targeted (green) regimens. **(b)** Tumor volumes following different regimens (color code as in a) versus untreated control (black curve). All treatments were administered continuously from day 5 to day 15 (gray shaded regions). **(c-f)** Phenotypic shifts of proliferation rate distribution *C*_*v*_ (*t, ρ*) under treatment. The proliferation rate *ρ* is shown on the x-axis, time in days is shown on the y-axis (blue = *t*_0_ = 0 days to yellow = *t*_*end*_ = 58 days). Please note that the y-axis scales differ. **(g-k)** Temporal heatmap of proliferation rate distribution *C*_*v*_ (*t, ρ*). (g) untreated, (h) uniform-targeting, (i) slow-targeting, (j) mid-targeting, and (k) high-targeting interventions. Density is encoded in colors from black (low density) to yellow (high density). Intensities of the treated groups are scaled to the highest observed density in the untreated group.

#### Low-proliferation targeting

Slowly proliferating cells experience maximal suppression, with efficacy decreasing exponentially as *ρ* increases. This selective pressure follows *f*_*low*_(*ρ*) = *ke*^*−λρ*^, where *λ* controls the decay rate of treatment efficacy along the proliferation axis (orange curve in Figure 4a).

#### Mid-proliferation targeting

Maximal suppression targets cells with intermediate proliferation rates via a Gaussian profile centered at the population’s initial mean proliferation rate 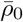. The treatment effect follows 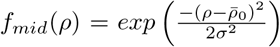 where *σ* determines the therapeutic breadth around the mean (purple curve in Figure 4a).

#### High-proliferation targeting

Rapidly proliferating cells are preferentially eliminated through an exponentially increasing treatment effect. The suppression profile is defined by *f*_*high*_(*ρ*) = *ke*^*λρ*^ where *λ* controls the growth rate of treatment efficacy (green curve in Figure 4a).

As expected, all treatment modalities reduced overall tumor burden relative to the untreated control (Figure 4b). However significant differences are observed in the primary clinical endpoints, as well as the phenotypic structure of the tumor.

Interestingly, this proliferation-structured model reproduces a characteristic feature of cytotoxic therapies such as chemotherapy and radiotherapy: tumor growth delay, a prolonged phase of growth suppression following the treatment window, before tumor regrowth resumes (Demidenko 2010; Yarnold et al. 1986). In classical unstructured growth models, this behavior typically requires the introduction of additional delay terms or compartments (e.g., quiescent or senescent cell states). In contrast, within our PDE framework, this phenomenon emerges naturally from the adaptive reshaping of the proliferation-rate distribution during and after treatment. Among all treatment strategies considered, selective targeting of intermediate proliferation phenotypes yields the greatest reduction in tumor burden and maximally prolongs the time to relapse (Figure 4b).

The temporal evolution of tumor structure under different treatment modalities was characterized by shifts in cell density across proliferation rates, as shown in Figure 4c-k. Density plots (Figure 4c-f) with time encoded in a color scale from blue (*t* = 0) to yellow (*t* = 58), and heatmaps (Figure 4g-k), with the proliferation rate on the x-axis, time on the y-axis, and density encoded in a color scale from black (low density) to yellow (high density), revealed progressive restructuring of phenotypic distributions, collectively depicting the temporal evolution of tumor cell dynamics under different treatment modalities.

Next to the inhibited growth during treatment, all treatments also changed the population composition with each treatment strategy inducing distinct compositional shifts (Figure 4c-f). Uniform targeting resulted in the elimination of cells across all phenotypes, leading to a decrease in density throughout the proliferation-rate spectrum (Figure 4c,h), narrowing the distribution compared to the untreated condition (Figure 4g). Slow-proliferation-targeted treatment caused a reduction in density within the low-proliferation-rate region, selecting for more rapidly proliferating cells and creating a noticeable shift to the right side of the proliferation rate distribution (Figure 4d,i). When targeting intermediate proliferation rates, the treatment gave rise to a bimodal distribution, where slow- and rapid-proliferating subpopulations coexisted, with very low densities in the mid-range (Figure 4e,j), a signature of an evolutionary branching process. Finally, targeting rapidly proliferating cells, a strategy that modulates the proliferation-mortality trade-off, led to a shift of the population toward lower proliferation rates during treatment (Figure 4f,k).

These phenotypic shifts are reflected in the dynamics of the population-wide mean proliferation rate. During the 10-day therapeutic window, all regimens drove a pronounced decline in 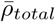 (Figure 5a), with the most substantial reduction observed under the mid-*ρ*-targeting treatment regime (purple points). The decline in the population-wide mean proliferation rate 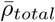 during treatment, even in slow-proliferation-targeted regimens, arose from the expanding doomed cell pool without proliferative capacity, reducing the population average. The pronounced decrease in 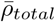 under the mid-proliferation-targeting regimen arises because this strategy targets the most densely populated region of the trait distribution, leading to the strongest depletion of viable cells and, consequently, a strong downward shift in the population-average proliferation rate.

**Fig. 5.**
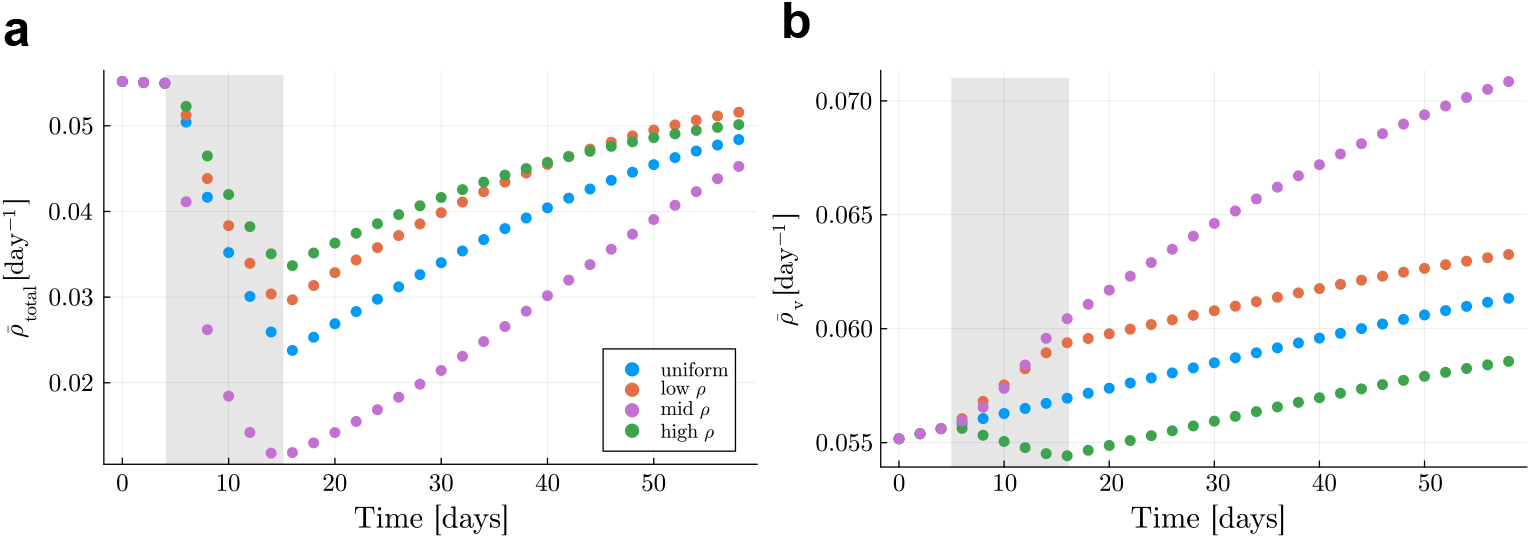
Mean proliferation rate dynamics across therapeutic regimens. **(a)** Population-wide mean proliferation rate 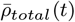. All regimens reduced 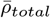 during treatment (days 5-15, gray shading), with mid-proliferation-targeted therapy (purple) showing the most pronounced decline. **(b)** Viable subpopulation mean proliferation rate 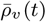. Uniform (blue), slow-targeted (orange), and mid-targeted (purple) regimens increased 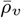 during treatment through selective elimination of slower-proliferating phenotypes, while high-targeted therapy (green) decreased 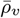 by preferentially eliminating rapidly proliferating clones. All groups exhibited a post-treatment increase in proliferation rates.

Post-treatment, the mean proliferation rate increased across all groups but did not fully reach the initial values observed before treatment, with the final values varying between treatment regimens.

When examining viable cells exclusively (Figure 5b), all treatment strategies except the high-targeting regime (green) increased 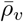 during the treatment window. However, the rate of change in 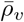 differed across treatment strategies. Under uniform targeting (blue), treatment did not alter the rate of increase in 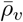. In contrast, targeting slowly proliferating cells (orange) accelerated the increase in 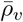 during treatment. The mid-proliferation-targeting regimen produced the strongest effect, yielding the highest rate of increase in 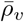 both during and after treatment, with a superlinear rise during the treatment window. By contrast, under high-proliferation targeting (green), 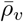 decreased during treatment because this strategy selectively depleted more rapidly proliferating cells and thereby favored slower-proliferating phenotypes. Immediately after treatment ended, the rate of change in 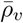 returned to its pre-treatment value for all strategies except mid-proliferation targeting, which maintained an elevated rate of increase.

Results of the model using the alternative trade-off term under treatment conditions are presented in Section S2.5.3.

## 4 Discussion

Our phenotype-structured PDE model reveals that intra-tumor heterogeneity in proliferation rate, driven by phenotypic diffusion and shaped by life-history trade-offs, fundamentally governs tumor growth dynamics and evolutionary trajectories. Model calibration to longitudinal data confirms that the trade-off is essential for accurate tumor volume prediction. Furthermore, we demonstrate that the maximum fitness strategy for proliferation rate dynamically declines as tumors grow, reflecting a shift toward slower, more resource-efficient phenotypes under increasing competition. Our *in silico* treatment experiments show that therapeutic interventions can reshape the proliferation landscape, leading to sustained effects on tumor growth dynamics beyond the treatment window. Moreover, selectively targeting different regions of the proliferation spectrum induces distinct patterns of phenotypic restructuring within the tumor population. In particular, our results indicate that a mid-proliferation targeting strategy, where the treatment profile overlaps with the most populated region of the proliferation distribution at the time of administration, maximizes the delay in tumor relapse. This contrasts with the conventional paradigm of preferentially targeting the fastest proliferating cells. In light of advances in targeted drug delivery systems (Cheng et al. 2023) and cell cycle–specific antitumor agents (Gross et al. 2023), these findings suggest a potential rationale for designing treatment strategies that exploit the underlying phenotypic distribution to achieve improved therapeutic efficacy.

A critical distinction from classical reaction-diffusion frameworks (e.g., Fisher-KPP equations) lies in our implementation of growth regulation. While Fisher-KPP systems typically employ *local* density dependence that supports traveling waves, our model imposes *global* resource competition through the logistic term where *N*(*t*) integrates all phenotypes. This global coupling enforces uniform nutrient constraints across the proliferation spectrum, reflecting the systemic resource limitations observed in solid tumors. Regarding phenotypic diffusion, we maintain a density-independent formulation. While density-dependent diffusion terms have been proposed in other contexts, such as tumor invasion (Tao and Tello 2015) and glioblastoma growth (Stepien et al. 2015) to capture spatial constraints, in our model diffusion represents genetic drift driven by stochastic mutations. Adopting density-dependent diffusion would imply that mutation rates scale with tumor size, a biologically unsupported premise.

Our model synergizes with the second-strike framework proposed by Gatenby et al. (2020), which proposes that tumors treated with a first treatment that reduces both tumor volume and ITH, become vulnerable to elimination by a second strike. While their model identifies the opportunity for extinction via first-strike-second-strike treatment, our framework provides the phenotypic roadmap: it predicts both the dominant traits and evolving heterogeneity patterns within the residual viable population after the first treatment. By predicting how heterogeneity shifts during treatment, we can select the second strike tailored to target the emergent phenotypic landscape. Thus, our contribution lies in offering a computational platform to identify and schedule synergistic treatment pairs where the first strike achieves sufficient volume reduction and the second strike selectively eliminates the resistant phenotype.

Furthermore, our results are consistent with findings from a compartmental ODE model of proliferation heterogeneity, which has primarily focused on the effects of targeting rapidly proliferating subpopulations and their impact on tumor growth and relapse dynamics (Raatz et al. 2021). A key advantage of incorporating heterogeneity in reproductive capacity as a continuously evolving trait within a PDE framework is that it enables a systematic exploration of diverse treatment modalities while naturally accommodating adaptive-dynamics principles within a classical tumor growth setting. This approach provides a mechanistic basis for the emergence of dominant clones and allows prediction of how their fitness evolves over time and in response to therapy. Taken together, these synergies between our framework and existing modeling approaches position the model as a tool for predicting relapse dynamics and designing combinatorial treatment strategies that leverage the tumor’s evolutionary trajectory, moving beyond monomorphic or compartmental descriptions toward a fully dynamic view of cancer evolution.

While our phenotype-structured PDE framework provides a powerful mechanistic description of proliferation-driven tumor dynamics, several aspects offer opportunities for future improvement. The model assumes a well-mixed, non-spatial tumor environment and does not explicitly account for spatial heterogeneity, such as nutrient gradients or physical barriers, which can strongly influence local selection pressures and clonal competition in solid tumors. Our model incorporates a single trade-off between proliferation and death, whereas in reality, phenotypic evolution is shaped by multiple, potentially interacting trade-offs involving key traits such as migration, metabolism, and therapy resistance. Extending the framework to include these additional dimensions would enable the representation of more complex adaptive strategies and a richer eco-evolutionary landscape.

Although our model accurately predicts macroscopic tumor growth, its predictions about the compositional shifts in phenotypic subpopulations, while mechanistically plausible, remain unvalidated due to limitations in longitudinal *in vivo* monitoring of phenotype distributions. Parameterization relies on fitting to bulk volume data and assumptions about initial phenotypic variance and trade-off structure, highlighting the need for more granular single-cell or lineage-tracing data to refine these estimates. Future work will extend the model to incorporate treatment data from clinical or preclinical trials, enabling direct comparison with observed resistance patterns. Moreover, for highly heterogeneous or spatially structured tumors, the PDE framework may be complemented by agent-based or hybrid models to capture emergent spatial and stochastic dynamics.

## 5 Acknowledgments

This work was Funded by the Deutsche Forschungsgemeinschaft (DFG, German Research Foundation) – GRK2762 – project number 450917483. We further acknowledge a travel grant provided by the GRK 2762 to L.S., which facilitated the initiation of the collaboration with the Russell Rockne’s Lab at City of Hope. We thank Haojun Chen and Xiaoyuan Chen for kindly providing the data associated with their publication (DOI: 10.1158/1535-7163.MCT-19-1098). We also thank Daniel Hoffmann and Tommaso Lorenzi for helpful discussions.

