## SupplementaryInformation for "Eco-Evolutionary Dynamics of Proliferation Heterogeneity: A Phenotype-Structured Model for Tumor Growth and Treatment Response"

### Supplementary Information

#### S1 Supplementary Materials and Methods

##### S1.1 Initial and Boundary Conditions

For computational implementation, the proliferation rate domain is truncated to $\rho \in [0, \rho_{\max}]$ where $\rho_{\max}$ is chosen such that $\rho_{\max} \gg \bar{\rho}(t) + 3\sigma(t)$ at all times to ensure that boundary conditions don't influence the solution. Homogeneous Neumann (zero-flux) boundary conditions are imposed at both ends of the domain:

$$\left.\frac{\partial C_v}{\partial \rho}\right|_{\rho=0} = \left.\frac{\partial C_v}{\partial \rho}\right|_{\rho=\rho_{\max}} = 0,$$

enforcing phenotypic conservation by preventing net gain or loss of cells through the boundaries.
The initial condition for viable cancer cells $C_v(t, \rho)$ is set as a normalized Gaussian distribution in proliferation rate space:

$$C_v(0, \rho) = \frac{N_{total,0}}{\sqrt{2\,\pi\,\sigma_0^2}}\, e^{-\frac{(\rho-\bar{\rho}_0)^2}{2\,\sigma_0^2}},$$

where $N_{total,0}$ is the initial tumor volume, $\bar{\rho}_0$ is the initial mean proliferation rate and $\sigma_0$ is the initial standard deviation. Given the small initial tumor volume, we assume no doomed cells are present at $t = 0$:

$$C_d(0, \rho) = 0 \quad \text{for all } \rho.$$

##### S1.2 Implementation and Simulation

All modeling and simulation workflows were performed in `Julia v1.12.0`. The partial differential equation system (Equations 1a-g) was implemented and discretized using `ModelingToolkit v10.26.0` (Ma et al. 2022) and `MethodOfLines v0.11.9` (MoL 2026) and solved using `OrdinaryDiffEq v.6.102.0` (Rackauckas and Nie 2017). Post-processing and visualization of time-resolved phenotype distributions, tumor volumes and mean proliferation rates were performed programmatically within the Julia ecosystem.

#### S1.3 Data

##### S1.3.1 Tumor Volume Data

To calibrate and validate our model, we used longitudinal tumor volume measurements from an untreated group of immunodeficient tumor-bearing mice reported in Zhao et al. (2020) comprising tumor growth trajectories of 11 replicates over 58 days post-implantation. To account for measurement errors due to caliper use, we smoothed the data using 1D splines with the `Dierckx v0.5.4` (Dierckx 2023) package in `Julia v1.12.0` (Bezanson et al. 2017) to ensure smooth and consistent tumor growth trajectories (see Section S1.3.2).

##### S1.3.2 Data preprocessing

Raw longitudinal tumor volume measurements obtained via calipers exhibit inherent experimental noise due to limitations in measurement precision, inter-operator variability, and challenges in consistently measuring irregular tumor geometries (Ogobuiro et al. 2025). To mitigate this noise and derive smoother, more consistent growth trajectories for model calibration and validation, we applied a one-dimensional cubic spline smoothing to the raw volume data (11 replicates over 58 days). This smoothing was implemented using the `Dierckx` package (v0.5.4) Dierckx (2023) within the `Julia v1.12.0` (Bezanson et al. 2017) programming environment. The primary objective was to reduce fluctuations attributable to measurement error while preserving the underlying biological growth trends. Appropriate smoothing parameters were selected to balance noise reduction with fidelity to the overall growth dynamics observed in the raw data to increase the robustness of model fitting.

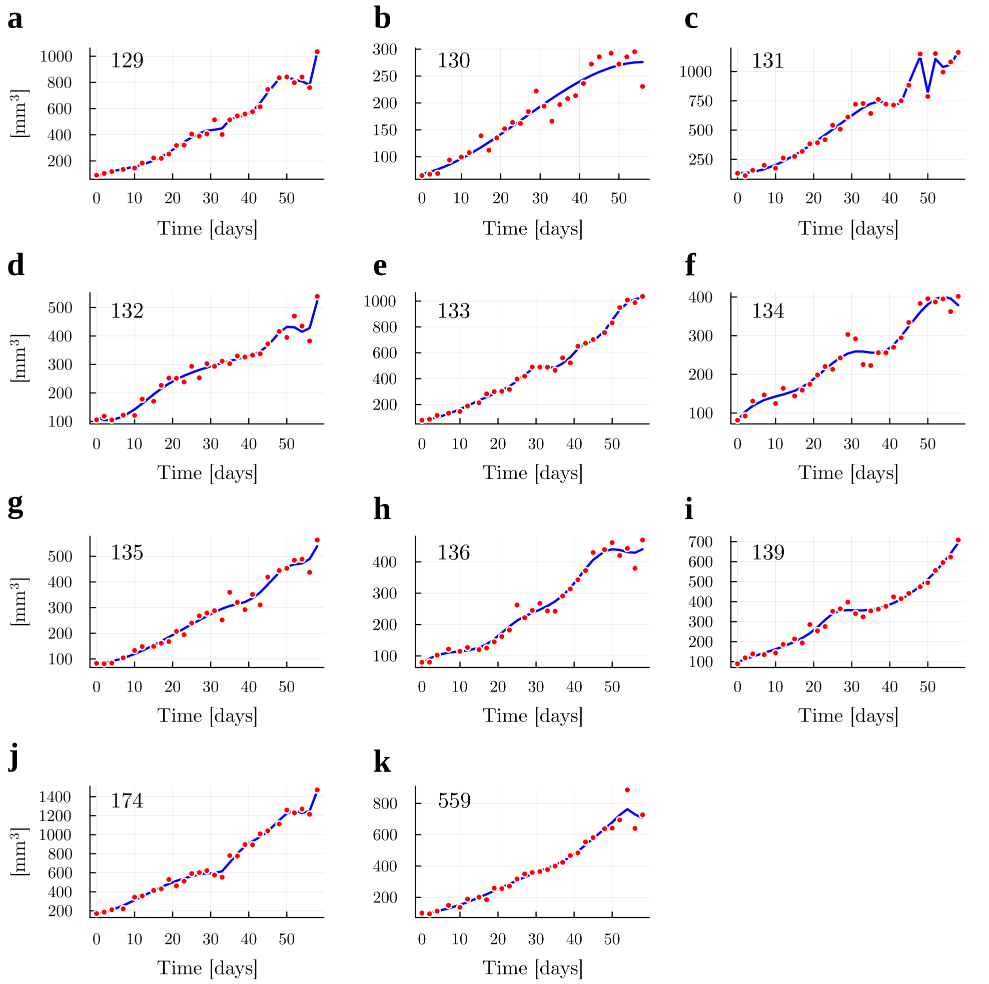


**Fig. S1** Raw tumor volume measurements and smoothed growth trajectories. Panels (a-k) display longitudinal data from 11 replicates over 58 days post-implantation. Red dots represent raw caliper measurements, exhibiting inherent experimental noise due to manual measurement variability and non-spherical tumor geometries. Blue curves show cubic spline fits applied to reduce non-biological fluctuations while preserving underlying growth trends. These smoothed trajectories were used for model calibration and validation.

Figure S1 presents the raw tumor volume measurements (red dots) alongside the smoothed growth trajectories (blue curves) for all 11 untreated control replicates (panels a-k) over 58 days post-implantation. The spline smoothing effectively reduced measurement noise inherent to caliper-based usage while preserving the underlying biological growth trends. However, in several replicates (notably panels a, c, j, k), substantial measurement fluctuations prevented complete noise elimination. In replicates like 129 and 174 (Figure S1a,j), a late-stage volume increase on day 58 remained present in the smoothed data. In replicate 131 (Figure S1c), pronounced fluctuations on days 48, 50, and 52 led to a persistent zig-zag pattern. In replicate 559 (Figure S1k), elevated tumor volume on day 54 followed by low volumes in days 56 and 58 led to a dip in the smoothed trajectory. In these cases, using higher smoothing parameters would have risked distorting biologically relevant features; we therefore prioritized retaining plausible growth dynamics over achieving perfect smoothness. Despite minor residual noise in these trajectories, the fitted curves capture the essential growth kinetics used for model calibration.

##### S1.4 Parameter Calibration

Model parameters were calibrated using longitudinal tumor volume data. The diffusion coefficient $D$, the initial standard deviation $\sigma_0$, the parameters governing the proliferation-mortality trade-off ($k_d$ and $s$), and the clearance rate for doomed cells $k_{cl}$ were fitted to replicate-level tumor growth trajectories in the PDE framework. The carrying capacity $K$ was individually calibrated per tumor replicate using an ODE-based logistic model to simplify PDE parameter estimation. Finally, the initial mean proliferation rate $\bar{\rho}_0$ was inferred via hierarchical Bayesian modeling across all 11 replicates' growth trajectories.
The simulation timeframe corresponds precisely to the experimental observation window (58 days), ensuring direct comparability with measured tumor dynamics.

#### S2 Supplementary Results

##### S2.1 Phenotypic variance and diffusion without life-history trade-offs amplify tumor growth

We performed comparative simulations to demonstrate the importance of proliferation-dependent life-history trade-offs for accurately capturing tumor dynamics (Figure S2a-b). PDE simulations with five distinct initial standard deviations ($\sigma_0$) omitting the trade-off term (Figure S2a, gradients of pink) exhibited accelerated growth compared to a homogeneous logistic growth ODE model of the form

$$\frac{dC}{dt} = \rho C \left(1 - \frac{C}{K}\right) \tag{S.1}$$

(Figure S2a, blue curve) at identical initial mean proliferation rates $\bar{\rho}_0$. The ODE model corresponds to the PDE model when $\sigma = 0$, where all cells share the same proliferation rate. In the ODE model as well as the PDE model without proliferation-mortality trade-off, the logistic term represents net growth (birth minus death). The diffusion coefficient $D$ was held constant across all simulations. We observed that the final tumor volumes scaled positively with initial phenotypic variance ($\sigma_0$), where wider initial distribution yielded larger tumors (Figure S2a, light pink curve) whereas narrower initial distributions yielded smaller tumors (Figure S2a, dark violet curve).

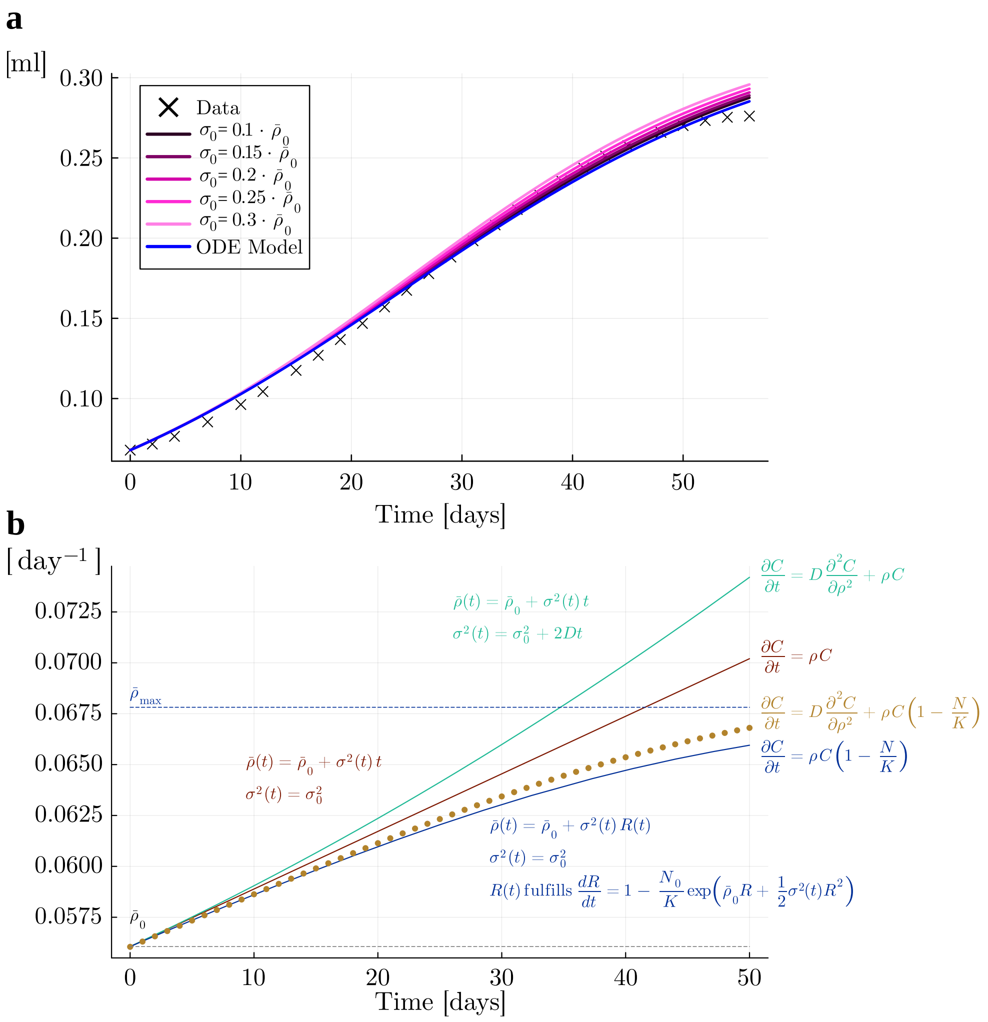


**Fig. S2** Comparative modeling simulations demonstrate the necessity of a proliferation-mortality trade-off. **(a)** Growth dynamics: PDE solutions without trade-off for increasing initial phenotypic variance (gradients of pink, dark=low variance, light=high variance) versus homogeneous ODE solution (blue curve) and experimental data (black crosses). **(b)** Mean proliferation rate dynamics: Exponential growth without diffusion (red), exponential growth with phenotypic diffusion (green), logistic growth without diffusion (dark blue), and logistic growth with phenotypic diffusion (yellow). The initial mean proliferation rate $\bar{\rho}_0$ is shown as a gray dotted line while the maximum proliferation rate $\bar{\rho}_{max}$ in the logistic growth case without diffusion is shown as a blue dotted line.

Mathematical analysis (Figure S2b) revealed mean proliferation rate dynamics $\bar{\rho}(t)$ across model variants, i.e. exponential and logistic growth, each with and without diffusion in phenotypic space. In exponential growth cases, the mean proliferation rate $\bar{\rho}(t)$ scales with phenotypic variance: $\bar{\rho}(t) = \bar{\rho}_0 + \sigma^2(t)\, t$. Without diffusion (red), the

variance remains constant ($\sigma^2(t) = \sigma_0^2$), leading to linear growth $\bar{\rho}(t) = \bar{\rho}_0 + \sigma_0^2\, t$. With diffusion (green), variance is amplified through diffusion ($\sigma^2(t) = \sigma_0^2 + 2Dt$), leading to superlinear growth $\bar{\rho}(t) = \bar{\rho}_0 + (\sigma_0^2 + 2Dt)\, t$. In the case of logistic growth without diffusion (blue), resource competition alters the dynamics of the mean proliferation rate which, in addition to the initial variance $\sigma_0^2$, depends on $R(t)$, i.e. $\bar{\rho}(t) = \bar{\rho}_0 + \sigma_0^2\, R(t)$, where $R(t)$ satisfies the saturating ODE $\frac{dR}{dt} = 1 - \frac{N_0}{K}\, e^{\bar{\rho}_0\, R + 0.5\sigma_0^2\, R^2}$. The exponential term in the ODE ($e^{\bar{\rho}_0\, R + 0.5\sigma_0^2\, R^2}$) represents cumulative growth potential, causing $R(t)$ and in consequence $\bar{\rho}(t)$ to saturate at $\bar{\rho}_{max}$ (Figure S2c, blue dotted line) when reaching carrying capacity $K$. For logistic growth with diffusion (yellow), numerical solutions showed persistent growth of the mean proliferation rate, with diffusion amplifying the effect of the variance on the mean growth rate. Derivations of these formulations are provided in the following sections (see Section S2.2).
In combination, these results demonstrate an inherent bias toward rapidly proliferating phenotypes that necessitates trade-off mortality to restore biological fidelity. This observation reflects the evolutionary interpretation of fitness as proliferative capacity, such that, in the absence of counterbalancing constraints, selection systematically favors increasingly fitter rapidly proliferating phenotypes.

#### S2.2 Derivation of mean proliferation rate dynamics

The mean proliferation rate $\bar{\rho}(t)$ is defined as the trait-weighted average over the tumor cell population (see Equation (S.2)). Since doomed cancer cells $C_d$ do not proliferate, this simplifies to a weighted average over the viable cell population $C_v(t, \rho)$:

$$\bar{\rho}(t) = \frac{\int_0^\infty \rho C_v(t,\rho) d\rho}{N(t)}\,, \tag{S.2}$$

where $N(t)$ is the tumor volume.

##### Exponential growth

We model viable cancer cell dynamics using the exponential growth PDE without diffusion in phenotypic space:

$$\frac{\partial C_v}{\partial t} = \rho\, C_v(t,\rho) \tag{S.3}$$

Solving Equation (S.3) yields:

$$C_v(t,\rho) = C_v(0,\rho)\, e^{\rho t}\,, \tag{S.4}$$

with initial condition $C_v(0, \rho)$ defined as a normal distribution centered at initial mean proliferation rate $\bar{\rho}_0$ with initial variance $\sigma_0^2$ and initial tumor volume $N_0$:

$$C_v(0,\rho) = \frac{N_0}{\sqrt{2\pi\sigma_0^2}}\, e^{-\frac{(\rho - \bar{\rho}_0)^2}{2\sigma_0^2}} \tag{S.5}$$

Substituting $C_v(0, \rho)$ in Equation (S.2) yields

$$\bar{\rho}(t) = \frac{\int \rho\, C_v(0,\rho)\, e^{\rho t}\, d\rho}{\int C_v(0,\rho)\, e^{\rho t}\, d\rho} = \frac{\int \rho \frac{N_0}{\sqrt{2\pi\sigma_0^2}}\, e^{-\frac{(\rho-\bar{\rho}_0)^2}{2\sigma_0^2}}\, e^{\rho t}\, d\rho}{\int \frac{N_0}{\sqrt{2\pi\sigma_0^2}}\, e^{-\frac{(\rho-\bar{\rho}_0)^2}{2\sigma_0^2}}\, e^{\rho t}\, d\rho}\,. \tag{S.6}$$

Combining exponents in Equation (S.6):

$$\bar{\rho}(t) = \frac{\int \rho\, e^{\rho t - \frac{(\rho-\bar{\rho}_0)^2}{2\sigma_0^2}}\, d\rho}{\int e^{\rho t - \frac{(\rho-\bar{\rho}_0)^2}{2\sigma_0^2}}\, d\rho} \tag{S.7}$$

Completing the square in the exponent yields:

$$\rho t - \frac{(\rho-\bar{\rho}_0)^2}{2\sigma_0^2} = -\frac{(\rho-\mu)^2}{2\sigma_0^2} + \bar{\rho}_0 t + \frac{\sigma_0^2 t^2}{2}, \quad \mu \equiv \bar{\rho}_0 + \sigma_0^2 t\,. \tag{S.8}$$

The integrals simplify using properties of Gaussian distributions. The denominator integrates to:

$$\int \exp\left(-\frac{(\rho-\mu)^2}{2\sigma_0^2}\right) d\rho = \sqrt{2\pi\sigma_0^2}\, \exp\left(-\bar{\rho}_0 t - \frac{\sigma_0^2 t^2}{2}\right)\,, \tag{S.9}$$

while the numerator gives:

$$\int \rho \exp\left(-\frac{(\rho-\mu)^2}{2\sigma_0^2}\right) d\rho = \mu\sqrt{2\pi\sigma_0^2}\, \exp\left(-\bar{\rho}_0 t - \frac{\sigma_0^2 t^2}{2}\right)\,. \tag{S.10}$$

Substituting (S.9) and (S.10) into (S.7) yields the linear relationship:

$$\bar{\rho}(t) = \bar{\rho}_0 + \sigma_0^2(t)\, t\,, \tag{S.11}$$

with

$$\sigma^2(t) = \sigma_0^2 \tag{S.12}$$

In the case of exponential growth with diffusion in phenotypic space, the variance increases linearly depending on the diffusion coefficient $D$, so that:

$$\sigma^2(t) = \sigma_0^2 + 2Dt\,, \tag{S.13}$$

leading to

$$\bar{\rho}(t) = \bar{\rho}_0 + \sigma_0^2(t)\, t \ = \bar{\rho}_0 + (\sigma_0^2 + 2Dt)t\,. \tag{S.14}$$

**Approximation Note:** This derivation assumes the trait distribution remains Gaussian. While the growth term $\rho\, C_v(t,\rho)$ introduces skewness and diffusion spreads the distribution, the Gaussian approximation accurately captures the first two moments ($\bar{\rho}(t)$ and $\sigma^2(t)$) for moderate variances and populations concentrated near the mean.

#### Logistic growth

Next, we derive the mean proliferation rate dynamics for the logistic growth case without diffusion in phentoypic space. The dynamics follow:

$$\frac{\partial C_v(t,\rho)}{\partial t} = \rho\, C_v(t,\rho)\left(1 - \frac{N(t)}{K}\right), \tag{S.15}$$

with total tumor volume $N(t)$ and carrying capacity $K$. For fixed $\rho$:

$$\frac{dC_v(t,\rho)}{dt} = \rho\, C_v(t,\rho)\left(1 - \frac{N(t)}{K}\right). \tag{S.16}$$

Solving via separation of variables:

$$\frac{1}{C}\frac{dC}{dt} = \rho\left(1 - \frac{N(t)}{K}\right) \implies \int_{C(0,\rho)}^{C(t,\rho)} \frac{dC}{C} = \rho \int_0^t \left(1 - \frac{N(s)}{K}\right) ds. \tag{S.17}$$

Defining

$$R(t) := \int_0^t \left(1 - \frac{N(s)}{K}\right) ds\,, \tag{S.18}$$

we obtain:

$$C_v(t,\rho) = C_v(0,\rho)e^{\rho R(t)}\,. \tag{S.19}$$

With Gaussian initial condition

$$C_v(0,\rho) = \frac{N_0}{\sqrt{2\pi\sigma_0^2}} \exp\left(-\frac{(\rho - \bar{\rho}_0)^2}{2\sigma_0^2}\right)\,, \tag{S.20}$$

Equation (S.19) becomes:

$$C_v(t,\rho) = \frac{N_0}{\sqrt{2\pi\sigma_0^2}} \exp\left(-\frac{(\rho - \bar{\rho}_0)^2}{2\sigma_0^2} + \rho R(t)\right)\,. \tag{S.21}$$

Completing the square in the exponent:

$$-\frac{(\rho - \bar{\rho}_0)^2}{2\sigma_0^2} + \rho R(t) = -\frac{(\rho - \bar{\rho}_0 - \sigma_0^2 R(t))^2}{2\sigma_0^2} + \bar{\rho}_0 R(t) + \frac{1}{2}\sigma_0^2 R(t)^2, \tag{S.22}$$

yields:

$$C_v(t,\rho) = \frac{N_0}{\sqrt{2\pi\sigma_0^2}} \exp\left(\bar{\rho}_0 R(t) + \frac{1}{2}\sigma_0^2 R(t)^2\right) \exp\left(-\frac{(\rho - \bar{\rho}(t))^2}{2\sigma_0^2}\right), \tag{S.23}$$

where

$$\bar{\rho}(t) = \bar{\rho}_0 + \sigma_0^2 R(t)\,, \tag{S.24}$$

is the mean proliferation rate. The total tumor volume $N(t)$ is computed as:

$$N(t) = \int_{-\infty}^{\infty} C_v(t,\rho)\,d\rho = N_0 \exp\left(\bar{\rho}_0 R(t) + \frac{1}{2}\sigma_0^2 R(t)^2\right). \tag{S.25}$$

Finally, $R(t)$ evolves as:

$$\frac{dR}{dt} = 1 - \frac{N(t)}{K} = 1 - \frac{N_0}{K}\exp\left(\bar{\rho}_0 R(t) + \frac{1}{2}\sigma_0^2 R(t)^2\right). \tag{S.26}$$

Thus, the mean proliferation rate follows the dynamics shown in Equation (S.24), with $R(t)$ fulfilling the saturating ODE in Equation (S.26). As $t \to \infty$, the population approaches carrying capacity, preventing further evolution in the absence of diffusion. Hence, $\bar{\rho}(t)$ saturates at a maximum value $\bar{\rho}_\infty$. To derive $\bar{\rho}_\infty$, we solve the system at steady state:

$$\frac{dR}{dt} \to 0 \quad \text{and} \quad N(t) \to K, \implies 1 = \frac{N_0}{K}\exp\left(\bar{\rho}_0 R_\infty + \frac{1}{2}\sigma_0^2 R_\infty^2\right), \tag{S.27}$$

The maximum mean proliferation rate is therefore:

$$\bar{\rho}_\infty = \bar{\rho}_0 0 + \sigma_0^2 R_\infty = \sqrt{\bar{\rho}_0^2 + 2\sigma_0^2 \ln(K/N_0)}\,, \tag{S.28}$$

showing how initial variance $\sigma_0^2$ and growth potential $(\ln(K/N_0))$ determine the population composition.

#### S2.3 Additional replicates for the monotonically increasing life-history trade-off term

We fitted the model to the tumor growth trajectories of ten replicates, not shown in the main text. These replicates were independently fitted using the same framework (See Section S1), with parameters calibrated to match experimental tumor volumes. The fitted parameter values can be found in Table S1 at the end of this section. Figure S3 shows tumor volume trajectories for these replicates. Despite experimental variability and noise, the model predictions (colored dots, with hue scaled by the population-wide mean proliferation rate $\bar{\rho}_{total}$) closely follow the experimental measurements (black crosses).

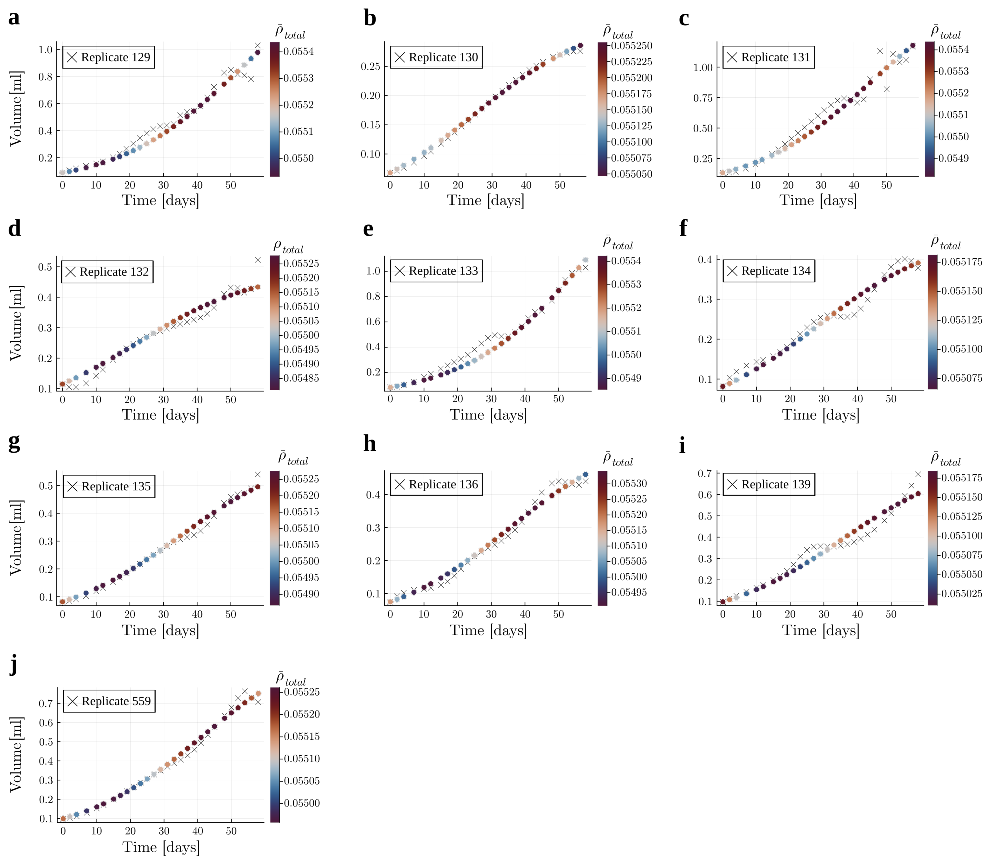


**Fig. S3** Tumor volume trajectories for the ten remaining replicates (beyond those in Figure 3a). PDE solutions (colored dots, hue scaled by population-wide mean proliferation rate $\bar{\rho}_{total}$ are plotted against experimental measurements (black crosses). Carrying capacity $K$ and parameters $D$, $\sigma_0^2$, $k_d$, $s$, and $k_{cl}$ were calibrated per replicate.

Figure S4 illustrates the spatiotemporal evolution of viable cell density across nine additional tumor replicates, showing the continuous restructuring of phenotypic composition throughout tumor progression. The distributions, visualized from $t = 0$ days (dark blue) to $t = 58$ days (yellow), evolve through diffusion-driven phenotypic variation, resulting in progressive broadening of the population-wide phenotype distribution. This spreading reflects the ongoing generation of phenotypic diversity, with both slow- and rapid-proliferating cells emerging over time. The population-wide mean proliferation rate $\bar{\rho}_{total}$ (red dots) remains stable over time, decreasing towards the end of the experimental observation window, which can be explained by the maximum fitness proliferation rate dynamics (see Section 3.2). The mean proliferation rate of the viable subpopulation, $\bar{\rho}_v$ (orange dots), shifts to the right, toward higher proliferation rates, over the experimental time window, indicating an enrichment of faster proliferating cells in the early stages, well below the carrying capacity. The initial phenotypic distributions are broadly similar across replicates. Two replicates (132 and 134) exhibit notably narrower initial distributions, suggesting reduced phenotypic heterogeneity at baseline. Despite these differences, the model captures the general trend of phenotypic diversification and selective enrichment of viable, high-proliferation phenotypes, highlighting the robustness of the evolutionary dynamics across biological variability.

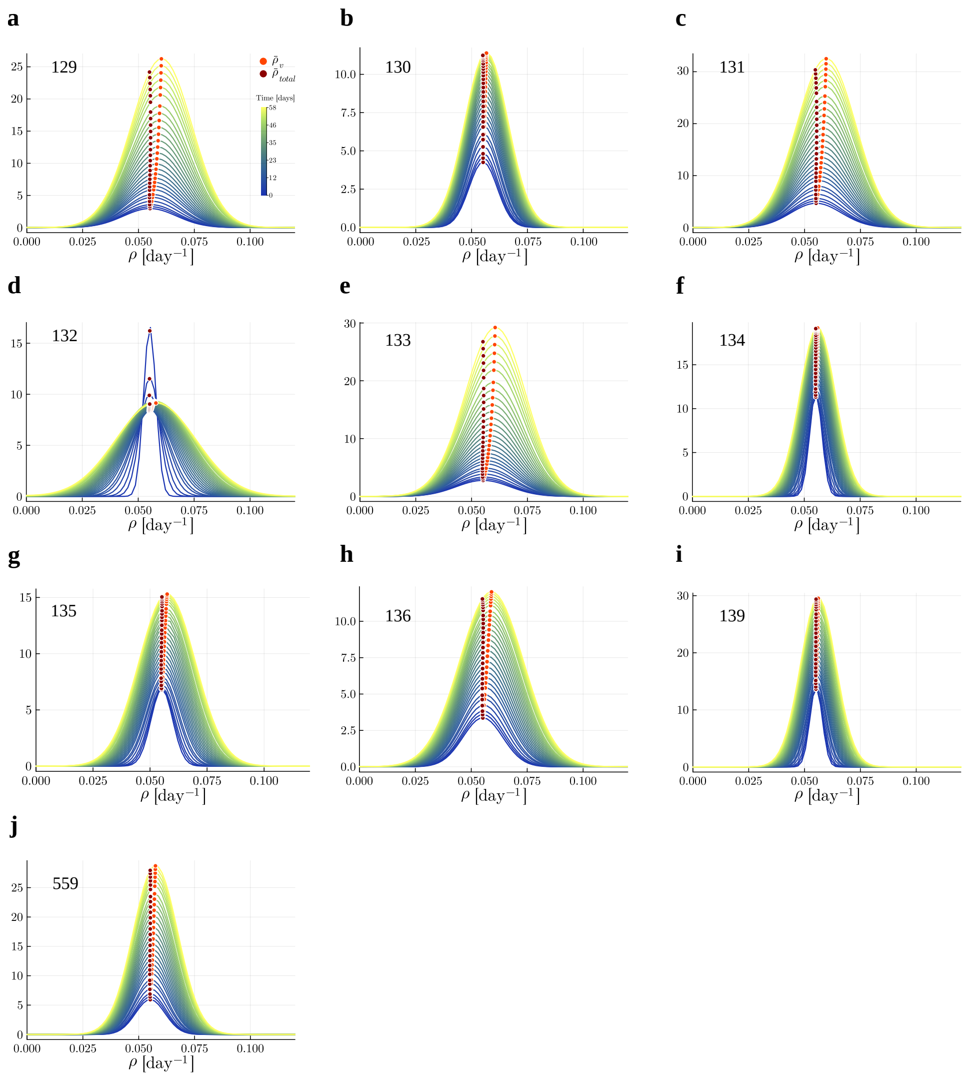


**Fig. S4** Viable cell densities for ten tumor replicates (beyond those shown in Figure 3b), shown from $t = 0$ days (dark blue) to $t_{end} = 58$ days (yellow). The population wide mean proliferation rate $\bar{\rho}_{total}$ is indicated as a red dot, the mean proliferation rate of the viable subpopulation $\bar{\rho}_v$ as an orange dot.

| Replicate ID | $k_d$ [day$^{(1-s)}$] | $s$ | $k_{cl}$ [day$^{-1}$] | $D$ [day$^{-3}$] | $\sigma_0$ [day$^{-1}$] |
|---|---|---|---|---|---|
| 559 | 0.07399 | 1.37088 | 0.001 | 4.86926E-07 | 0.00662 |
| 129 | 106.53624 | 3.69561 | 0.001 | 4.86926E-07 | 0.01189 |
| 130 | 0.17998 | 1.8718 | 0.00134 | 4.86926E-07 | 0.00633 |
| 131 | 32.03829 | 3.30708 | 0.001 | 4.86926E-07 | 0.01131 |
| 132 | 0.02309 | 1 | 0.001 | 2.7335E-06 | 0.00276 |
| 133 | 212.80353 | 3.92203 | 0.001 | 4.86926E-07 | 0.01203 |
| 134 | 0.00781 | 1 | 0.001 | 4.86926E-07 | 0.00276 |
| 135 | 0.02287 | 1 | 0.001 | 1.13326E-06 | 0.00474 |
| 136 | 0.04599 | 1.05253 | 0.001 | 1.06869E-06 | 0.00888 |
| 139 | 0.01012 | 1 | 0.001 | 4.86926E-07 | 0.00276 |
| 174 | 16.13731 | 3.26268 | 0.001 | 4.86926E-07 | 0.0069 |

**Table S1** Fitted values for each experimental replicate for the monotonically increasing trade-off term for the parameters $k_d$, $s$, $k_{cl}$, $D$, and $\sigma_0$ with their respective units.

#### S2.4 Fitness landscape for the monotonically increasing life-history trade-off term

The net growth function $g(\rho, N)$ balances reproductive advantages against survival costs, creating a fitness peak, maximizing fitness in the current microenvironment. For an exemplary tumor with the parameters $N_0 = 10\,\text{mm}^3$, $K = 1000\,\text{mm}^3$, $k_d = 1.5$ days, and $s = 2.0$ and a fixed ratio of tumor burden to carrying capacity ($\frac{N}{K} = 0.7$, corresponding to $t = 47$ days), the net growth rate $g(\rho)$ (Figure S5a, black curve) exhibits a maximum at $\rho^* \approx 0.1$ (Figure S5a, red point), dividing the fitness landscape into two areas: In the growth-dominated region ($\rho < \rho^*$, Figure S5a, yellow shaded area), increasing proliferation increases fitness as reproductive gains outweigh survival costs. Conversely, in the death-dominated area ($\rho > \rho^*$, Figure S5a, gray shaded area), an increase in proliferation rate reduces fitness as survival costs outweigh growth potential.

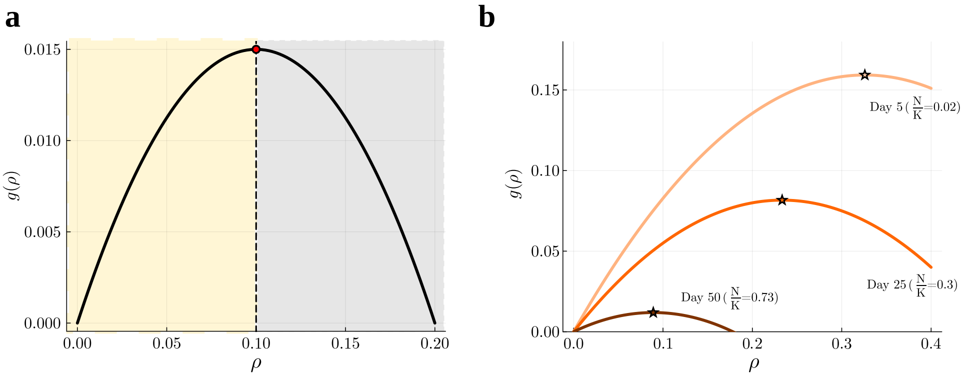


**Fig. S5** Fitness landscape in an exemplary tumor **(a)** Fitness landscape at $\frac{N}{K} = 0.7$ for an exemplary tumor with the parameters $N_0 = 10\,\text{mm}^3$, $K = 1000\,\text{mm}^3$, $k_d = 1.5$ days, and $s = 2.0$. The net growth rate $g(\rho)$ exhibits a maximum at $\rho^* \approx 0.1\,\text{day}^{-1}$ (red point). For $\rho < \rho^*$, the growth potential $\rho\left(1 - \frac{N}{K}\right)$ dominates (yellow shaded area), while death $k_d\rho^s$ dominates for $\rho > \rho^*$. **b** The net growth rate $g(\rho)$ and $\rho^*$ are reshaped progressiveley during tumor progression, examples are given for day 5 ($\frac{N}{K} = 0.02$, light orange curve), day 25 ($\frac{N}{K} = 0.3$, dark orange curve), and day 50 ($\frac{N}{K} = 0.73$, brown curve). Stars mark $\rho^*$ which is shifted leftward toward slower proliferation rates as the tumor approaches carrying capacity.

The net growth rate $g(\rho)$ and its evolutionary stable maximum, $\rho^*$, are dynamically coupled to tumor volume $N(t)$ (see Figure 2b). This induces a shift in the fitness landscape as the tumor grows: early-stage tumors ($N \ll K$, light orange curve in Figure S5b) have higher net growth rates than larger tumors (dark orange and brown curves in Figure S5b). Specifically, $\rho^*$ decreases from 0.33 [$\text{day}^{-1}$] on day 5 ($\frac{N}{K} = 0.02$, light orange star in Figure S5b) to $\rho^* = 0.23\,[\text{day}^{-1}]$ on day 25 ($\frac{N}{K} = 0.3$, dark orange star in Figure S5d) and to $\rho^* = 0.09\,[\text{day}^{-1}]$ on day 50 ($\frac{N}{K} = 0.73$, brown star in Figure S5b). Early stage tumors favor high proliferation phenotypes to use abundant resources, but increasing mortality penalties at high $\rho$ constrain this strategy, in favor of low proliferative cells that can cope the more competitive environment better. Consequently, $\rho^*$ declines as $N(t)$ grows (see Equation (3)), dark orange and brown curves in Figure S5d), reflecting selection toward slower phenotypes under saturation. This implies that tumors continuously evolve along a trajectory of decelerating proliferation as tumor burden increases.

#### S2.5 Analysis for an alternative trade-off term

We implemented an alternative trade-off term $d(\rho)$, that punishes cells with very low and very high proliferation rates:

$$d(\rho) = k_d \frac{(\rho - \rho_{opt})^2}{(\rho - \rho_{opt})^2 + \rho\,(\rho_{max} - \rho)} \tag{S.29}$$

The aim of this analysis was to investigate the impact of evolutionary constraints on cells that transition into a state of permanent arrest or markedly reduced proliferative capacity. We performed similar analyses to the ones shown in the main text for the monotonically increasing trade-off term. The simulation results incorporating the U-shaped trade-off term indicated that, although the trade-off shape does not alter the macroscopic tumor growth trajectory, it has a substantial impact on microscopic phenotypic restructuring.

##### S2.5.1 Model fits

The PDE model (Equations 1a-g) was fitted to longitudinal tumor volume measurements from 11 tumor replicates (see Section S1.3) through optimization of parameters $k_d$, $\rho_0$, $k_{cl}$, $D$, and $\sigma_0$ for each replicate. Carrying capacity $K$ was individually calibrated per tumor replicate using an ODE-based logistic model to simplify PDE parameter estimation.

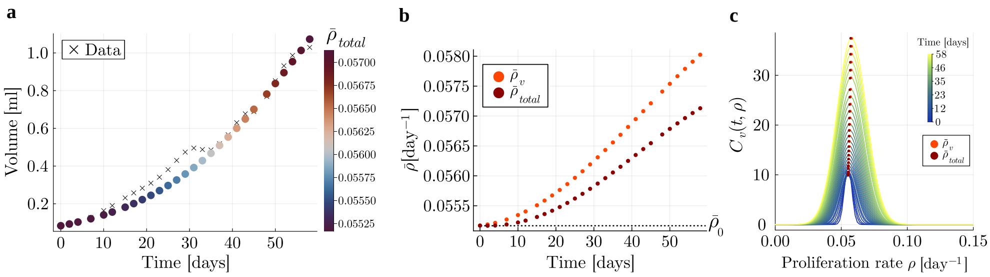


**Fig. S6** Model validation and phenotypic dynamics for tumor replicate 131 **(a)** Tumor volume trajectories: PDE solutions (colored dots, hue scaled by population-wide mean proliferation rate $\bar{\rho}_{total}$) versus experimental measurements (black crosses). **(b)** Proliferation rate evolution: Temporal dynamics of $\bar{\rho}_{total}$ (entire population, red) and $\bar{\rho}_v$ (viable subpopulation, orange). Black dotted line indicates initial $\bar{\rho}_{total,0} = 0.05516$ [day$^{-1}$]. **(c)** Phenotypic restructuring: Viable cell distributions $C_v(t,\rho)$ from $t_0 = 0$ days (dark blue) to $t_{end} = 58$ days (yellow). Red and orange dots mark $\bar{\rho}_{total}(t)$ and $\bar{\rho}_v(t)$ at sampled time points. Parameters $\rho_0 = 0.054$ [day$^{-1}$], $k_d = 0.170$ [day$^{-1}$], $k_{cl} = 0.130$ [day$^{-1}$], $D = 1.149 \times 10^{-6}$ [day$^{-3}$], $\sigma_0 = 0.00276$ [day$^{-1}$] were fitted; carrying capacity $K = 2.25$ [ml] was replicate-specific.

The model solutions are visualized for a representative replicate (Figure S6). The results for the remaining replicates are provided in the next section (see Section S2.6). The predicted growth trajectories show close agreement with the experimental measurements (Figure S6a), similar to the results for the monotonically increasing trade-off term shown in the main text (see Section 3.3). It captured the logistic growth pattern, while simultaneously resolving the underlying phenotypic evolution.

Temporal patterns in the population-wide mean proliferation rate $\bar{\rho}_{total}$ (color-coded) were observed during the experimental period, with detailed dynamics illustrated in Figure S6b. Unlike the monotonically increasing trade-off, the mean proliferation rate of the entire population $\bar{\rho}_{total}$ (red) and of the viable subpopulation $\bar{\rho}_v$ (orange) increased from day 0 to the end of the experimental observation period.

A continuous restructuring of the population composition occurred throughout tumor progression, as visualized in Figure S6c. The initial Gaussian profile (dark blue) increased in amplitude, representing an increase in tumor volume, and broadened progressively due to diffusion-driven phenotypic variation. The parameter values for all replicates are shown in Table S2 in Section S2.6.

##### S2.5.2 Long-term behavior

For the trade-off term presented in Equation (S.29), we simulated the model for a tumor with the parameters $K = 1.77$ [ml], $\rho_0 = 0.053$ [day$^{-1}$], $k_d = 0.128$ [day$^{-1}$], $k_{cl} = 0.083$ [day$^{-1}$], $D = 1.644 \times 10^{-5}$ [day$^{-3}$], $\sigma_0 = 0.00276$ [day$^{-1}$] for a time period of 2000 days. The total tumor volume $N_{total}(t)$, consisting of the volumes of viable cells $N_v$ and doomed cells $N_d$, initially grew toward carrying capacity and stabilized at carrying capacity (Figure S7a) after 150 days. During tumor growth, the viable subpopulation grew (orange curve in Figure S7b) until it reached a steady state in a similar time frame to the total tumor volume. The population of doomed cells followed a similar trajectory, characterized by initial growth followed by a stabilization at a constant volume.

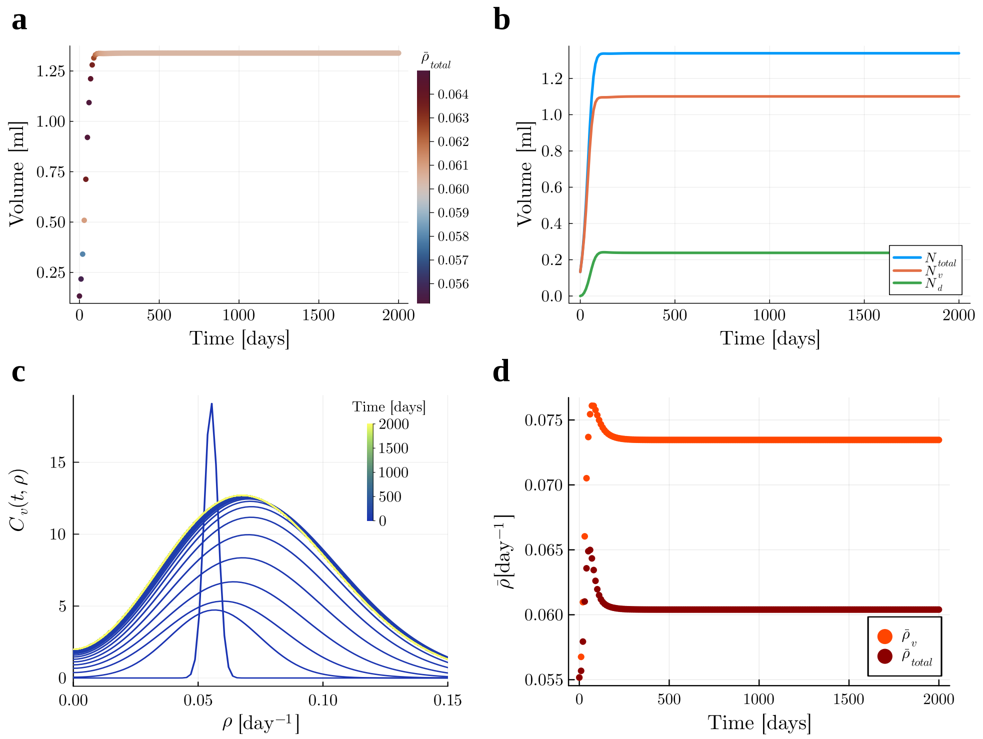


**Fig. S7** Long-term model behavior for a tumor with the parameters $K = 1.77$ [ml] $\rho_0 = 0.053$ [day$^{-1}$], $k_d = 0.128$ [day$^{-1}$], $k_{cl} = 0.083$ [day$^{-1}$], $D = 1.644 \times 10^{-5}$ [day$^{-3}$], $\sigma_0 = 0.00276$ [day$^{-1}$] for a time period of 2000 days. **(a)** Tumor volume trajectories: PDE solutions for the total tumor volume $N_{total}(t)$ (colored dots, hue scaled by population-wide mean proliferation rate $\bar{\rho}_{total}$). **(b)** Trajectories of the total tumor volume $N_{total}$ (blue curve), the volume of the viable subpopulation $N_v$ (orange curve), and the volume of the doomed subpopulation $N_d$ (blue curve). **(c)** Phenotypic restructuring: Viable cell distributions $C_v(t, \rho)$ from $t_0 = 0$ days (dark blue) to $t_{end} = 2000$ days (yellow). **(d)** Proliferation rate evolution: Temporal dynamics of $\bar{\rho}_{total}$ (entire population, red) and $\bar{\rho}_v$ (viable subpopulation, orange).

This accelerated stabilization in comparison to the monotonically increasing trade-off term leads to a fast and stable phenotypic restructuring (Figure S7c). During tumor growth, the initial distribution (dark blue) grows and broadens, and rapidly converges to a stable profile (yellow). Correspondingly, the mean proliferation rates $\bar{\rho}_{total}$ and $\bar{\rho}_v$ increase during tumor growth, then decline to steady states within the same time frame (Figure S7d). Notably, this trade-off induces an interior stable strategy, whereby the phenotype distribution converges to a non-trivial fixed point located away from the phenotype boundaries.

##### S2.5.3 Treatment explorations for U-shaped trade-off term

Similar to the analysis in Section 3.4 in the main text, we investigate the effects of four different proliferation-rate-dependent treatment strategies on the tumor volumes and population composition of an exemplary tumor.

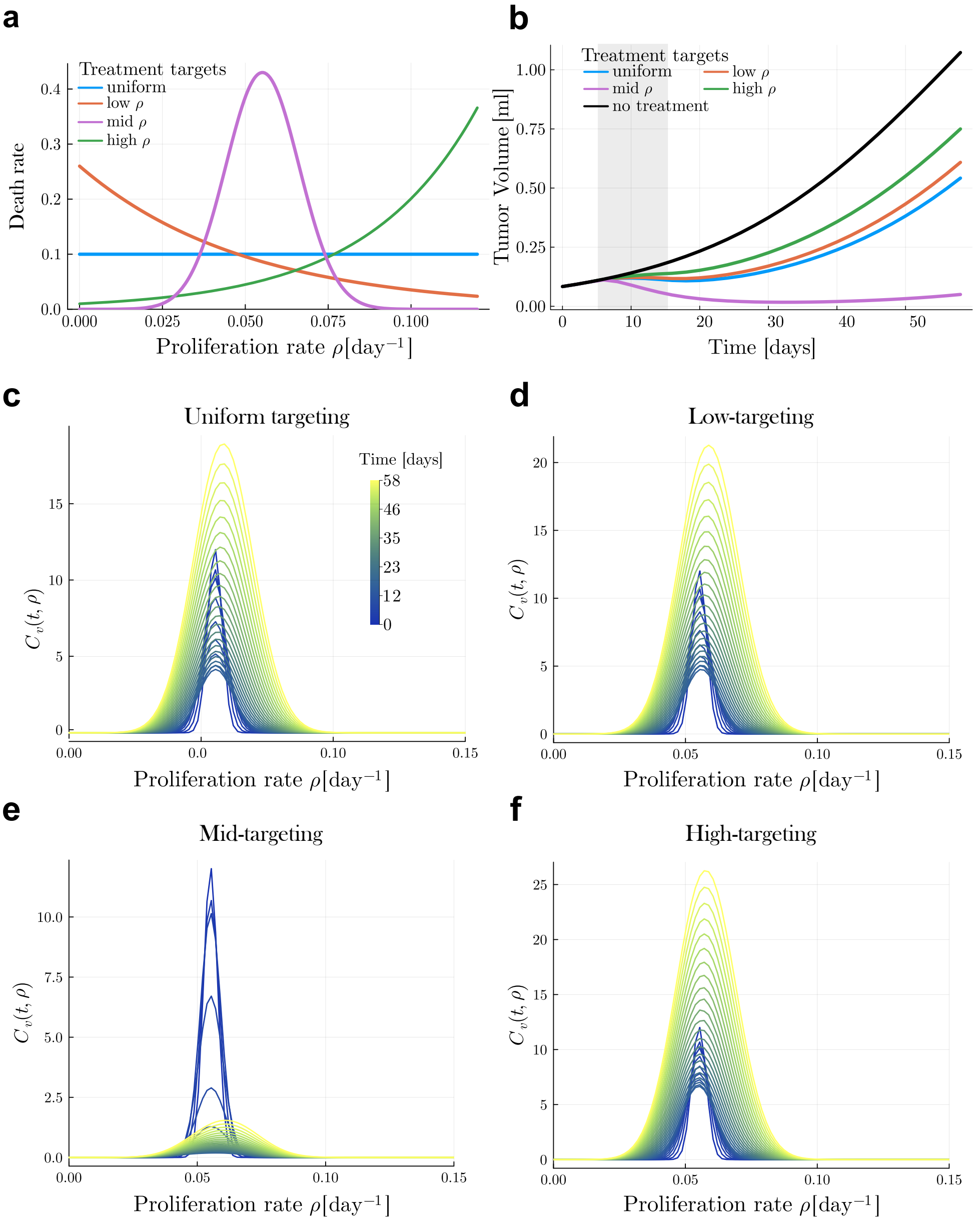


**Fig. S8** Treatment-induced restructuring of phenotypic landscapes using the U-shaped proliferation-mortality trade-off. **(a)** Therapeutic targeting profiles: Suppression functions for uniform (blue), slow-proliferation-targeted (orange), mid-proliferation-targeted (purple), and high-proliferation-targeted (green) regimens. **(b)** Tumor volumes following uniform-targeting (blue curve), low-proliferation targeted (orange curve), mid-proliferation targeted (purple curve), and high-proliferation targeted (green curve) regimes versus untreated control (black curve). All treatments were administered continuously from day 5 to day 15 (gray shaded regions). **(c-f)** Phenotypic shifts under treatment: Density plots of $C_v(t, \rho)$ for (b) uniform, (c) slow-targeting, (d) mid-targeting, and (e) high-targeting interventions.

The tumor volumes using the four different treatment strategies (Figure S8a) are shown in Figure S8b. Following all treatments, the tumor burden remained lower than in the untreated group (black curve). During the mid-proliferation targeting treatment, the tumor volume decreases (purple curve), followed by a stagnation until day 50 and an increase at the end of the simulation time frame. Treatment with the low-proliferation (orange curve) and uniform (blue curve) treatments led to a slight decrease in tumor volume followed by an increase about 15 days after treatment finished. The high-proliferation treatment led to a constant tumor volume during treatment and a regrowth shortly thereafter (green curve). The macroscopic tumor growth trajectories under different treatment strategies remain comparable across the two trade-off formulations.
The phenotypic shifts shown in Figure S8c-f are similar to the ones following treatments in the model with the monotonically increasing trade-off term: Following the uniform treatment (Figure S8c), the distribution is shifted towards higher proliferation, with the population's peak residing leftward of the pre-treatment distribution's peak (blue curves). A similar pattern can be observed under the low-proliferation targeting treatment (Figure S8d). Following the mid-proliferation targeting treatment, the density decreases dramatically during treatment, leading to a narrow distribution that only slowly regrows. The density is almost depleted after treatment and slowly grows back after treatment (Figure S8e). Following the high-proliferation targeting, the distribution is pushed down. The mean proliferation rate stays almost constant (Figure S8f). Following treatment, the distributions broaden.

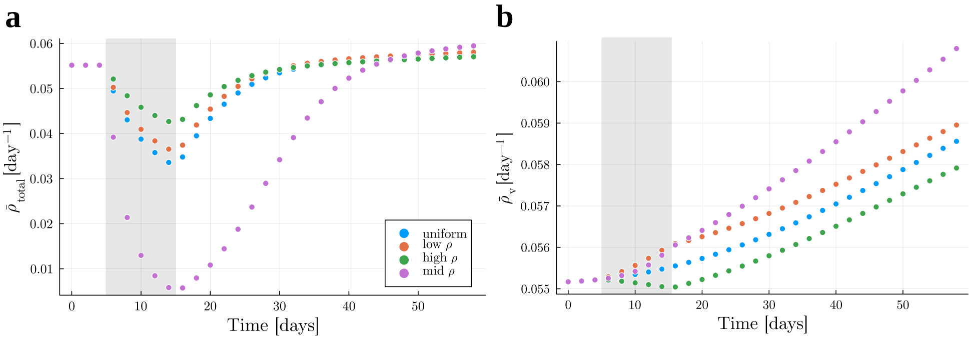


**Fig. S9** Divergent proliferation dynamics and fitness landscape remodeling across therapeutic regimens. **(a)** Population-wide mean proliferation rate $\bar{\rho}_{total}(t)$. All regimens reduced $\bar{\rho}_{total}$ during treatment (days 5-15, gray shading), with mid-proliferation-targeted therapy (violet) showing the most pronounced decline. **(b)** Viable subpopulation mean proliferation rate $\bar{\rho}_v(t)$. Uniform (blue), slow-targeted (orange), and mid-targeted (violet) regimens increased $\bar{\rho}_v$ during treatment through selective elimination of slower-proliferating phenotypes, while high-targeted therapy (green) decreased $\bar{\rho}_v$ by preferentially eliminating fast-proliferating clones. All groups exhibited post-treatment increase in proliferation rates.

The same can be observed in the mean proliferation rates after treatment (Figure S9). The population-wide mean proliferation rate $\bar{\rho}_{total}$ decreased in response to all treatments (Figure S9a). The strongest decline in $\bar{\rho}_{total}$ can be observed following the mid-proliferation targeting treatment (purple). After all treatments, $\bar{\rho}_{total}$ increased to a value close to the initial value. The viable subpopulation's mean proliferation rate $\bar{\rho}_v$ increased following the uniform (blue), low-targeting (orange), and mid-targeting (purple) treatments (Figure S9b). Following the high-targeting treatment $\bar{\rho}_v$ decreased slightly and increased again after the treatment was over (green). The overall pattern of phenotypic restructuring following treatment was comparable between the two examined trade-off functions, with a more accelerated dynamics reaching steady state in the U-shaped trade-off.

#### S2.6 Additional replicates for U-shaped trade-off term

We fitted the model using the U-shaped trade-off term to the tumor growth trajectories of ten replicates not shown in Section S2.6. These replicates were independently fitted using the same framework, with parameters calibrated to match experimental tumor volumes. Figure S10 shows tumor volume trajectories for these replicates. The model predictions (colored dots, with hue scaled by the population-wide mean proliferation rate $\bar{\rho}_{total}$) closely follow the experimental measurements (black crosses).

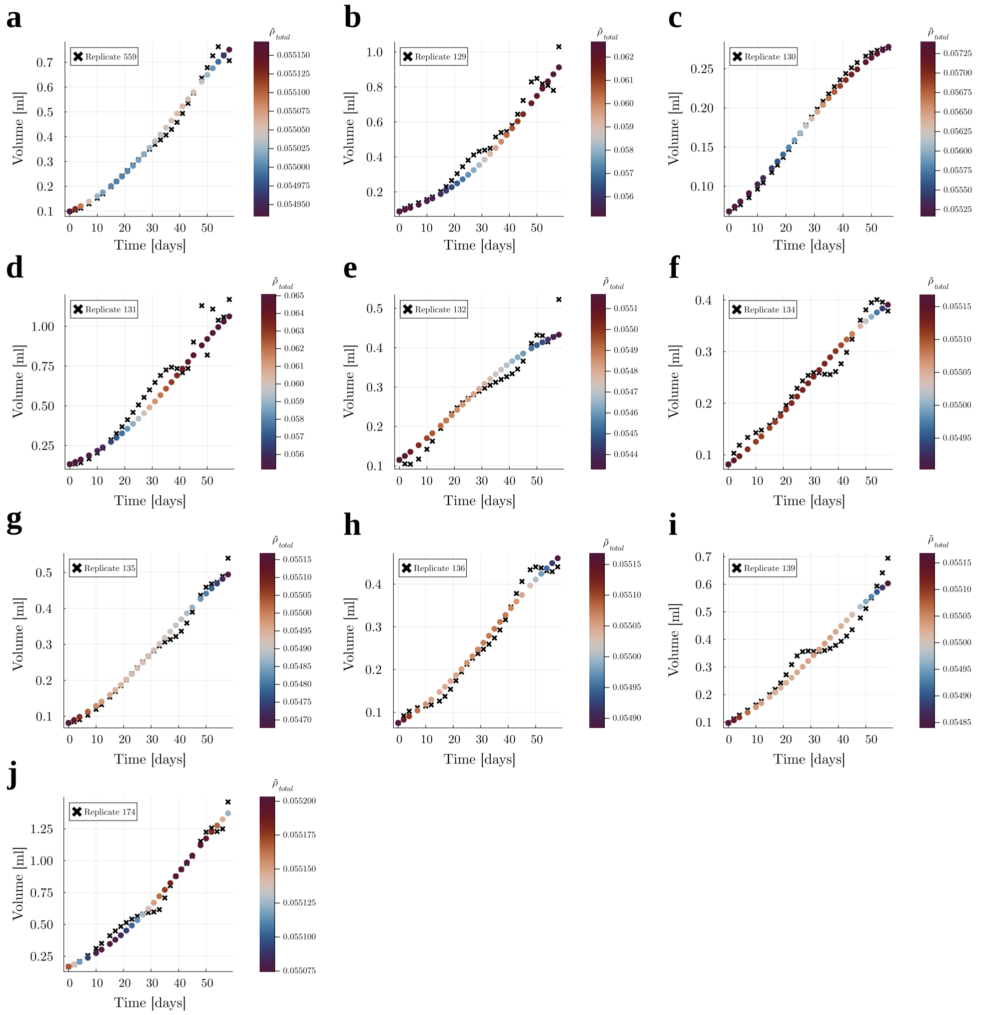


**Fig. S10** Tumor volume trajectories for the ten remaining replicates (beyond those in Figure S6a,d). PDE solutions (colored dots, hue scaled by population-wide mean proliferation rate $\bar{\rho}_{total}$ are plotted against experimental measurements (black crosses). Carrying capacity $K$ and parameters $D$, $\sigma_0^2$, $k_d$, $\rho_0$, and $k_{cl}$ were calibrated per replicate.

The fitted tumor growth trajectories are close to the data, similar to what we fitted with the monotonically increasing proliferation-mortality trade-off term.

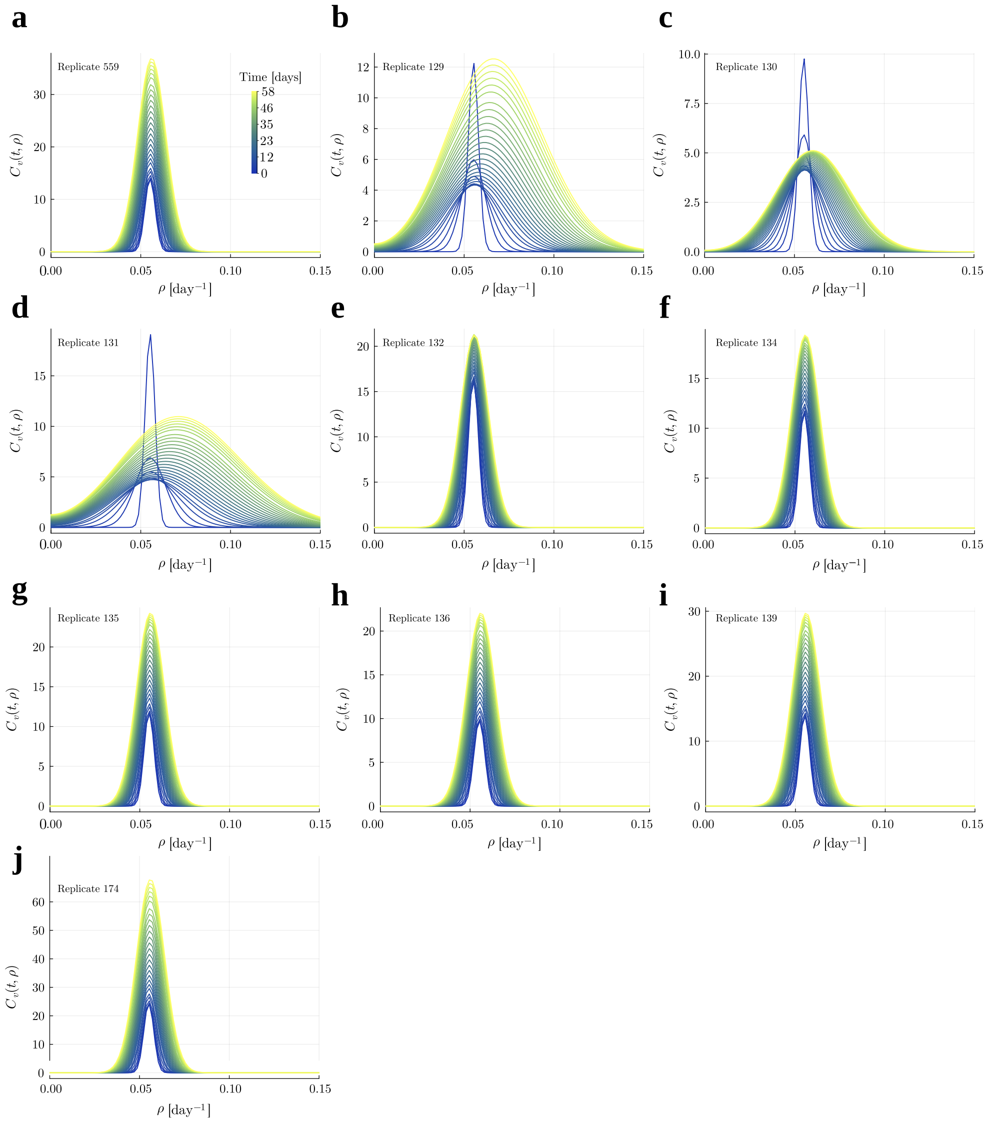


**Fig. S11** Viable cell densities for nine tumor replicates (beyond those shown in Figure S6b,e), shown from $t = 0$ days (dark blue) to $t_{end} = 58$ days (yellow).

Figure S11 illustrates the spatiotemporal evolution of viable cell density across nine additional tumor replicates, showing the continuous restructuring of phenotypic composition throughout tumor progression. The distributions, visualized from $t = 0$ days (dark blue) to $t = 58$ days (yellow), evolve through diffusion-driven phenotypic variation, resulting in progressive broadening of the population-wide phenotype

| Replicate ID | $k_d$ [day$^{-1}$] | $\rho_{opt}$ [day$^{-1}$] | $k_{cl}$ [day$^{-1}$] | $D$ [day$^{-3}$] | $\sigma_0$ [day$^{-1}$] |
|---|---|---|---|---|---|
| 559 | 0.04315 | 0.04548 | 0.001 | 4.86926E-07 | 0.0028211 |
| 129 | 0.13495 | 0.05173 | 0.09088 | 8.76448E-06 | 0.0028615 |
| 130 | 0.09398 | 0.05838 | 0.04534 | 4.4408E-06 | 0.0027584 |
| 131 | 0.12824 | 0.05315 | 0.08323 | 1.64409E-05 | 0.0027584 |
| 132 | 0.01917 | 0.03924 | 0.001 | 4.86926E-07 | 0.0027584 |
| 133 | 0.16964 | 0.05432 | 0.13006 | 1.14931E-06 | 0.0027584 |
| 134 | 0.04559 | 0.0484 | 0.001 | 4.86926E-07 | 0.0027584 |
| 135 | 0.03337 | 0.04337 | 0.001 | 4.86926E-07 | 0.0027584 |
| 136 | 0.0422 | 0.04611 | 0.001 | 5.04269E-07 | 0.0030524 |
| 139 | 0.04201 | 0.04599 | 0.001 | 4.86926E-07 | 0.0027584 |
| 174 | 0.05449 | 0.04791 | 0.001 | 4.86926E-07 | 0.0027584 |

**Table S2** Fitted values for each experimental replicate for the monotonically increasing trade-off term for the parameters $k_d$, $\rho_{opt}$, $k_{cl}$, $D$, and $\sigma_0$ with their respective units.

distribution. This spreading reflects the ongoing generation of phenotypic diversity, with both slow- and fast-proliferating cells emerging over time. In comparison to the model fits using the monotonically increasing trade-off, the U-shaped trade-off leads to narrower initial distributions.

#### S2.7 Trade-off structure shapes proliferation distributions and dynamics

The monotonically increasing trade-off term (Equation (1a)) penalizes rapid-growing cells exclusively, whereas the U-shaped trade-off term (Equation (S.29)) imposes a symmetric penalty on both very rapid- and slow-growing cells. This difference in penalty structure directly shapes the model dynamics. In the absence of treatment, the U-shaped trade-off inherently favors intermediate proliferation rates, resulting in a narrower initial proliferation rate distribution compared with the broader distribution observed under the monotonically increasing trade-off. Steady states in $N_{total}$, $N_v$, and $N_d$ are reached more rapidly with the U-shaped trade-off, reducing the time required for phenotypic restructuring.

The resulting steady-state distributions also differ. Under the monotonically increasing trade-off, the distribution becomes flattened and skewed toward lower proliferation rates, whereas the U-shaped trade-off produces a more symmetric distribution with a high density around intermediate rates. These differences in final population composition directly reflect the distinct selective pressures imposed by the two trade-off mechanisms. Despite these microscopic differences, both trade-offs produce similar tumor growth curves.

Under treatment, the behaviors of the two models are largely comparable. In both cases, the population-wide mean proliferation rate $\bar{\rho}_{total}$ decreases during treatment and increases again afterward. With the U-shaped trade-off, $\bar{\rho}_{total}$ returns to its initial value within the simulated period of 58 days, whereas it remains below the initial value when the monotonically increasing trade-off is used. This difference reflects the faster convergence to steady state under the U-shaped trade-off. The viable population's mean proliferation rate, $\bar{\rho}_v$, shows similar trends under both trade-offs: it increases during treatment for the uniform, low-targeting, and mid-targeting treatments, but decreases under the high-targeting treatment.
